# Polypyrimidine Tract-Binding Protein 1 and hnRNP A1 recognize unique features of the Sm site in U7 snRNA

**DOI:** 10.1101/2023.08.19.553944

**Authors:** Xiao-cui Yang, Zbigniew Dominski

## Abstract

U7 snRNA is a 60-nucleotide component of U7 snRNP, a multi-subunit endonuclease that cleaves precursors of metazoan replication-dependent histone mRNAs at the 3’ end, hence generating mature histone mRNAs. The Sm site in U7 snRNA differs from the Sm site in spliceosomal snRNAs and promotes the assembly of a unique Sm ring containing Lsm10 and Lsm11 instead of SmD1 and SmD2 found in the spliceosomal snRNPs. The assembly of the spliceosomal Sm ring depends on the SMN complex, with one of its nine subunits, Gemin5, recognizing the spliceosomal Sm site. While the assembly of the U7-specific Sm ring also requires the SMN complex, the unusual Sm site of U7 snRNA is not recognized by Gemin5, and the identity of its counterpart that performs this function in biogenesis of U7 snRNP, has not been determined. Here, we looked for proteins that bind U7 snRNA but not to its mutant altered within the Sm site. We identified Polypyrimidine Tract-Binding Protein 1 (PTBP1) as the main protein that meets this specificity. Binding of PTBP1 to U7 snRNA also depends on the upstream CUCUUU motif that base pairs with histone pre-mRNAs and defines substrate specificity of U7 snRNP. Thus, PTBP1 simultaneously recognizes two functionally essential and highly conserved sites within U7 snRNA. In addition to PTBP1, U7 snRNA interacts with hnRNP A1, which recognizes a different part of the U7-specific Sm site. Interestingly, the two proteins can form with U7 snRNA a larger complex, which also contains SMN protein, a subunit of the SMN complex. Altogether, these results raise the possibility that PTBP1 and hnRNP A1 act collectively to substitute for Gemin5 in the assembly of U7-specific Sm ring.

## INTRODUCTION

In metazoans, 3’ end processing of replication-dependent histone pre-mRNAs does not follow the two-step pathway of cleavage coupled to polyadenylation and instead involves single-step cleavage, yielding mature histone mRNAs terminated with a highly conserved stem-loop structure followed by a 5-nucleotide single stranded tail (1). The cleavage reaction is conducted by U7 snRNP, a multi-component RNA-guided endonuclease consisting of U7 snRNA and several proteins, including CPSF73, which functions as the catalytic component (2–4). The same role is played by CPSF73 in the canonical cleavage/polyadenylation machinery (5,6).

The substrate specificity of U7 snRNP is dictated by U7 snRNA, an approximately 60-nucleotide RNA whose 5’ region base pairs with a purine rich region known as Histone pre-mRNA Downstream Element (HDE) in histone pre-mRNAs (7,8). Centrally located within U7 snRNA is an Sm binding site of 11 nucleotides with the AAUUUGUCUAG consensus that nucleates the assembly of a ring composed of seven proteins: Lsm10, Lsm11, SmE, SmF, SmG, SmB and SmD3 (9,10). This sequence differs from the 9-nucleotide spliceosomal consensus, AAUUUUGG, which promotes the assembly of a canonical Sm ring containing Sm proteins D1 and D2 instead of Lsm10 and Lsm11. Lsm10 and Lsm11 are largely responsible for the role of U7 snRNP as the endonuclease in 3’ end processing of histone pre-mRNAs. Lsm11, the largest protein of the Sm/Lsm family, interacts with FLASH, a protein of 220 kDa, and these two U7 snRNP subunits act together to recruit the cleavage module consisting of symplekin, CPSF100 and CPSF73 (11,12). Both Lsm11 and Lsm10 make additional contacts with this module, helping to stabilize and arrange the complex and regulate its nucleolytic activity (3).

The assembly of the canonical heptameric ring on spliceosomal snRNPs occurs through a highly controlled and multi-step process carried out by two large entities: PRMT5 methylosome and SMN complex (13–17). The PRMT5 methylosome consists of three components (PRMT5, Mep50 and pICln) and acts to pre-organize the seven Sm subunits into smaller sub-complexes, SmD1/D2, SmE/F/G and SmB/D3 (18,19), and to symmetrically dimethylate C-terminal arginines in three Sm proteins: D1, B and D3 (20–22). The SMN complex consists of 9 components (SMN, Gemin2-8 and Unrip) and brings together each spliceosomal snRNA and the three Sm sub-complexes, promoting a stepwise assembly of a complete ring that surrounds the Sm site. Within the SMN complex, Gemin2 binds a transient pentamer consisting of the Sm subunits D1, D2, E, F and G (23). Gemin5 recognizes the Sm site, allowing ring assembly only on spliceosomal snRNAs, preventing illegitimate interactions of Sm proteins with random RNA sequences (24–26). SMN binds symmetrically demethylated arginines on the Sm subunits D1, B and D3, facilitating an orderly transfer of the preassembled Sm sub-complexes onto the Sm site and displacement of Gemin5 (27–30). How exactly this occurs and what role is played by the remaining SMN subunits remain largely unknown. Mutations in the SMN gene cause spinal muscular atrophy (SMA), a genetic disorder characterized by selective death of motor neurons and muscle degeneration (16,31–36). SMA is believed to result from inefficient biogenesis of spliceosomal snRNPs and perturbations in alternative splicing events important for motor neuron function and maintenance (37–39). Alternatively, SMN may perform a tissue-specific function in motor neurons, explaining selective degeneration of only these cells (40–42).

The assembly of the unique Sm ring of U7 snRNP is also under the control of the PRMT5 methylosome and the SMN complex (19,43). *In vitro*, PRMT5 methylates arginines in the N-terminal regions of Lsm11 and SmE (44), and this unusual methylation pattern may serve as a specificity determinant for the assembly of U7 snRNP, helping to distinguish this pathway from the canonical and much more ubiquitous mechanism that operates during the biogenesis of spliceosomal snRNPs. The importance of the SMN complex in the biogenesis of the U7 snRNP is best illustrated by the observation that RNA-mediated downregulation of the SMN protein results in a gradual decrease of the amount of U7 snRNP, accumulation of longer U7 snRNA precursors and a defect in 3’ end processing of histone pre-mRNAs (45,46). Consistent with the distinct composition of the U7 ring, the Sm site of U7 snRNA is not recognized by Gemin5 (24), which is likely replaced by another RNA binding protein that ensures the recruitment of Lsm10 and Lsm11, preventing potentially harmful incorporation of SmD1 and SmD2. It remained unclear whether this task is carried out by a component of the SMN complex different than Gemin5, or by an unrelated protein, which subsequently delivers U7 snRNA to this complex for the completion of the assembly process.

Here, we carried out biochemical experiments to identify proteins in mammalian cell extracts that bind wild type U7 snRNA but fail to bind mutant U7 snRNA whose Sm site was converted to the spliceosomal consensus. Our studies identified Polypyrimidine Tract-Binding Protein 1 (PTBP1) (47–49) as the main protein displaying this binding specificity. The interaction of PTBP1 with U7 snRNA was abolished not only by changing the Sm site, but also by mutations within the highly conserved pyrimidine motif that base pairs with histone pre-mRNAs and facilitates the recruitment of substrates to U7 snRNP. Thus, PTBP1 alone can simultaneously recognize two features of the U7 snRNA critically important for the function of the U7 snRNP in 3’end processing. PTBP1 contains four RNA Recognition Motifs (RRMs) that have high affinity for CU-containing sequences (50–52), the hallmark of both the substrate base pairing element and Sm site in U7 snRNA. In addition to PTBP1, U7 snRNA also interacts with hnRNP A1 (53,54). This protein recognizes the UAG trinucleotide (55,56), a different characteristic feature of the U7-specific Sm site conspicuously absent from the spliceosomal Sm sites. Interestingly, hnRNP A1 and PTBP1 can form a larger complex with U7 snRNA and SMN, potentially linking this complex to U7 snRNP assembly.

PTBP1 uses its RRM2 to interact with a group of proteins that share the (S/G)(I/L)LGxxP consensus motif, including Matrin3, Raver1, Raver2, CCAR1 and CCAR2 (57–61). Why these proteins, often containing their own RNA binding domains, form an intimate complex with PTBP1 is not clear but they likely act by assisting PTBP1 in its multiple functions, including alternative splicing (57,62). Surprisingly, U7 snRNA stabilizes complexes of PTBP1 with these proteins, while an unrelated RNA containing multiple CU dinucleotides and tightly binding PTBP1 acts in the opposite manner, weakening the same complexes. This observation may be indicative of U7 snRNA being involved in processes other than U7 snRNP biogenesis.

Both PTBP1 and hnRNP A1 are versatile RNA binding proteins with links to multiple processes, including regulation of alternative splicing, mRNA stability, 3’ end processing and cap-independent translation, and they have multiple paralogs in vertebrates (58,63) (54). In human cells PTBP1 has two paralogues, PTBP2 and PTBP3, that share as much as 75% sequence identity that likely play both redundant and specific roles (64,65). PTBP2 is mostly expressed in neuronal cells and is one of the key players in differentiating neuronal progenitor cells into mature neurons (66–69). U7 snRNA interacts with all three PTBP paralogues and one attractive possibility is that this and possibly other specific components of U7 snRNP become repurposed for new roles in terminally differentiated, post-mitotic cells that cease to replicate DNA and express replication-dependent histone proteins.

## RESULTS

### Human proteins that bind U7 snRNA

Mature human U7 snRNA consists of 60-63 nucleotides and three functional/structural regions. The sequence of nucleotides 11-18 is highly conserved among distant metazoan species and have the CUCUUU consensus. We refer to this sequence as the Substrate Base Pairing (SBP) site (Fig. 1A). The SBP site forms a duplex with Histone pre-mRNA Downstream Element (HDE), a purine-rich sequence in histone pre-mRNA located downstream of the site cleaved by the U7 snRNP (8,70). The formation of this duplex is the primary mechanism that defines substrate specificity of U7 snRNP. Nucleotides 21-31 encompass the Sm site. This region mediates the assembly of an Sm ring of U7 snRNP consisting of Lsm10, Lsm11 and five proteins shared with the spliceosomal snRNPs: SmB, SmD3, SmE, SmF and SmG (9). The 3’ half of U7 snRNA folds into an extended stem-loop structure. U7 snRNA is synthesized by RNA polymerase II as a longer precursor, which undergoes co-transcriptional cleavage by the Integrator complex (71,72), yielding a longer U7 snRNA species extended at the 3’ end by a single stranded tail of different length. This extended tail is removed in the cytoplasm following the assembly of a complete Sm ring on U7 snRNA (73).

**Fig. 1.**
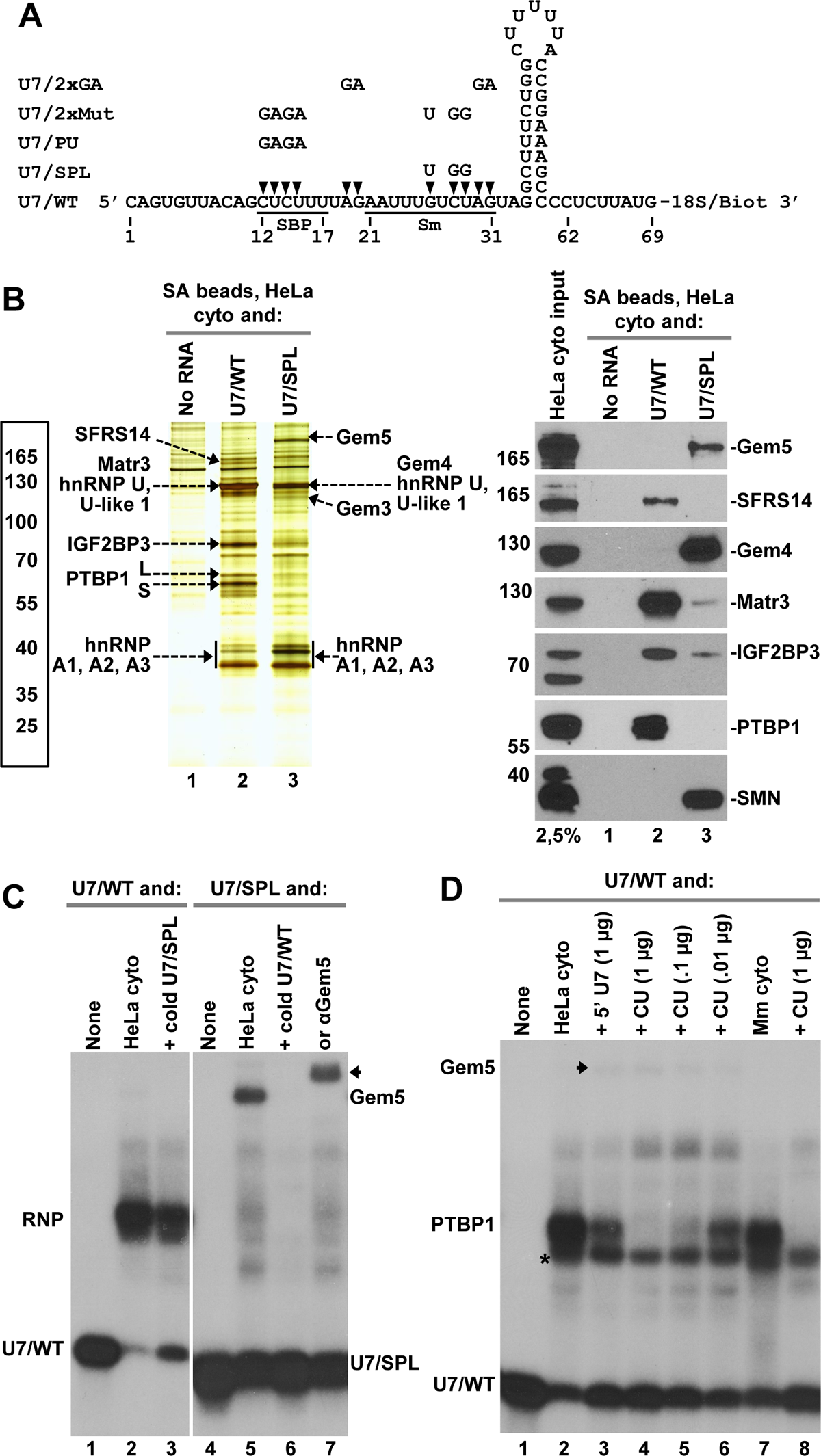
Human proteins that bind U7 snRNA. **A.** The sequence and secondary structure of the 69-nucleotide U7/WT RNA containing biotin and an 18-atom spacer (18S) at the 3’ end. The first 63 nucleotides correspond to mature U7 snRNA found in U7 snRNP. The last 6 nucleotides are present in U7 snRNA precursor and *in vivo* are deleted following the Sm ring assembly. The Substrate Base Pairing (SBP) site between nucleotides 12-17, and the Sm site between nucleotides 21-31 are underlined. Nucleotides indicated with arrowheads were substituted with the sequences shown above, giving rise to U7 snRNA mutants indicated to the left. **B.** Affinity purification of proteins bound to U7 snRNA on streptavidin (SA) beads via 3’ biotin. A HeLa cytoplasmic extract was incubated without RNA (lane 1) or in the presence of U7/WT or U7/SPL RNAs and bound proteins were immobilized on SA beads, separated in a 4-12% SDS/polyacrylamide gel and detected by silver staining (left) or Western blotting (right). Protein bands identified by mass spectrometry are indicated. In the right panel, the input lane contains ∼2.5% of the cytoplasmic extract used in the binding experiment. **C and D.** Electrophoretic mobility shift assay (EMSA) to detect formation of RNA/protein complexes. U7/WT and U7/SPL RNAs were labeled at the 5’ end with ^32^P, mixed with a cytoplasmic extract alone or in the presence of RNA competitors, as indicated, and separated in a 6% native polyacrylamide gel. The shift in the position of the radioactive probes upon forming protein complexes was visualized by autoradiography. Probes alone are shown in lanes 1 and 4 (panel C) and in lane 1 of panel D. With the exception of lanes 7 and 8 of panel D, which contain a cytoplasmic extract from mouse myeloma cells, all lanes contain a HeLa cytoplasmic extract. In lane 7 of panel C, the sample contains antibody against Gemin5.

One of the most outstanding questions in the biogenesis of U7 snRNP is how the specific Sm site of U7 snRNA is recognized, ultimately resulting in the incorporation of Lsm10/11 heterodimer and preventing an erroneous incorporation of the more abundant SmD1/D2 heterodimer that occupies the same position in the spliceosomal ring. To identify proteins that recognize the U7-specific Sm site and may act as the functional counterpart of Gemin5, we incubated U7 snRNA with cell extracts and immobilized bound proteins on streptavidin beads via biotin attached to U7 snRNA either cis or trans, as previously described (74,75). Among multiple experimental variations tested (length of U7 snRNA and methods of its synthesis, position of biotin, incubation conditions and protocols for extract preparation), the most consentient results were obtained with a chemically synthesized 69-nucleotide U7 snRNA containing wild type Sm site (AAUUUGUCUAG) and an extended single stranded tail and biotin at the 3’ end (Fig. 1A). Compared to the 63-nucleotide mature U7 snRNA that is part of active U7 snRNP (3), the 69-nucleotide RNA is extended at the 3’ end by 6 nucleotides (Fig. 1A), hence mimicking the U7 snRNA precursor (U7 pre-snRNA), which is co-transcriptionally generated by the Integrator complex in the nucleus. In the cytoplasm, this precursor serves as the genuine substrate for the assembly of the Sm ring and prior to returning to the nucleus with the complete ring is trimmed at the 3’ end to 60-63 nucleotides (73).

In parallel to the 69-nucleotide RNA referred to as U7/WT, we used U7/SPL RNA whose Sm site was altered by substituting three nucleotides to match the spliceosomal consensus of AAUUU<u>u</u>U<u>gg</u>AG (nucleotide substitutions are indicated with small letters), as previously described (76,77). U7/WT and U7/SPL RNAs were incubated at 4 °C with HeLa cytoplasmic extracts and bound proteins were immobilized on streptavidin (SA) beads via the 3’ biotin tag and separated by SDS/PAGE. To select high affinity binders, the amount of RNA in this experiment was limited to 750 ng for 750 µl of the cytoplasmic extract. Silver staining visualized three major protein bands in the material bound to U7/WT RNA, but not to U7/SPL RNA (Fig. 1B, left panel, lanes 2 and 3, respectively). Two of these bands migrating at around 60 kDa were identified by mass spectrometry as two splicing isoforms, long (L) and short (S), of Polypyrimidine Tract-Binding Protein 1 (PTBP1) (47–49). Both isoforms contain four RNA Recognition Motifs (RRMs), with the long isoform likely containing an additional 26-residue segment that separates RRM2 and RRM3. The third unique protein band migrating above the 70 kDa size marker was identified as Insulin-Like Growth Factor 2 mRNA-Binding Protein 3 (IGF2BP3). The identity of PTBP1 and IGF2BP3 was confirmed by Western blotting (Fig. 1B, right panel, lane 2).

The U7/SPL mutant RNA, while failing to bind PTBP1 and IGF2BP3, interacted with a unique protein migrating at about 185 kDa, identified by mass spectrometry and Western blotting and as Gemin5 (Fig. 1B, left and right panels, lane 3). The presence of Gemin5 only in the material purified via the mutant U7 snRNA but not via the WT U7 snRNA confirms previous reports based on using different methods that this component of the SMN complex readily interacts with the spliceosomal-type Sm site but has no affinity for the U7-specific Sm site (24), validating our *in vitro* approach. To our knowledge, this is the first direct illustration of endogenous Gemin5 discriminating between the two types of snRNAs and a clear indication that the recognition of the Sm site in U7 snRNA is mediated by a different protein. In addition to Gemin5, the material purified by U7/SPL contained other components of the SMN complex, including Gemin3, Gemin4 and SMN (Fig. 1B, left and right panels, lane 3). We note that the presence of Gemin3 and Gemin4 was detected in only some extracts. All these proteins are conspicuously absent from the material bound by U7/WT RNA (Fig. 1B, left and right panels, lane 1).

Among other proteins identified by mass spectrometry, the material purified via U7/WT RNA contained Matrin3 (Matr3), a known binding partner of PTBP1 and SFRS14 (Fig. 1B, right panel, lane 1), a splicing factor that likely co-purifies with Matrin3 (58). Both RNAs interacted with two highly abundant proteins, hnRNP U and U-like 1, that co-migrate with Matrin3 and in silver-stained gels masked the enrichment of Matrin3 by U7/WT RNA. In addition to hnRNP U and U-like 1, U7/WT and U7/SPL RNAs also interacted with three highly related shorter hnRNPs: A1, A2 and A3 (see below).

To confirm these results by a different method and to demonstrate that PTBP1 directly binds U7/WT RNA, we carried out electrophoretic mobility shift assays (EMSA). U7/WT and U7/SPL RNAs were labeled at the 5’ end with ^32^P, mixed with a cytoplasmic extract, and RNA/protein complexes were separated in a 6% native polyacrylamide gel. The assay revealed a striking difference in the ability of each RNA to form RNP complexes (Fig. 1C). U7/WT probe migrates in native polyacrylamide gels near the bottom of the gel and in the presence of a cytoplasmic extract was shifted to a higher position near the middle of the gel due to forming a larger complex (Fig. 1C, lanes 1 and 2). The specific shift was not eliminated by a 500-fold molar excess of U7/SPL unlabeled RNA (Fig. 1C, lane 3). Thus, the protein that binds to U7/WT has much lower affinity for U7 snRNA mutant containing a spliceosomal-type Sm site. To determine whether the shift is due to binding PTBP1, IGF2BP3 or Matrin3, we used specific antibodies against each of these three proteins. A small fraction of the complex was super-shifted to a higher position only by an antibody against PTBP1, with other antibodies having no effect (not shown). One interpretation of this result is that the complex is formed by PTBP1, but the incomplete super-shift was due to low accessibility of the anti-PTBP1 antibody to PTBP1 in the complex. Repeated EMSA experiments failed to detect in native gels larger complexes that could potentially contain PTBP bound to both U7 snRNA and Matrin3. It is possible that these complexes do not survive electrophoretic separation.

When 5’-labeled U7/SPL RNA was used as a probe, the characteristic shift seen for U7/WT RNA was not observed and instead a different complex with a lower mobility was formed (Fig. 1C, lanes 4 and 5). The complex was quantitatively super-shifted by an antibody against Gemin5 (Fig. 1C, lane 7, arrow), identifying the bound protein as Gemin5, consistent with the results of pull-down experiments. A 500-fold molar excess of U7/WT cold RNA abolished this interaction (Fig. 1C, lane 6), suggesting that the U7-specific Sm site has a detectable affinity for Gemin5. Note that both Gemin5 and PTBP1 are abundant proteins, and they both can be sufficiently enriched for detection in silver-stained gels.

PTBP1 tightly interacts with RNAs containing CU-rich regions (50,51,78,79). In the presence of the CU oligonucleotide (5’ CUCUCUCUCUUUCUUU 3’) containing multiple PTBP1 binding sits, the formation of the main U7/WT complex was inhibited (Fig. 1D, lanes 4-6), supporting the notion that the interacting protein is indeed PTBP1. Note that the use of this competitor revealed the presence of a different U7/WT RNA/protein complex that was insensitive to the excess of this oligonucleotide and migrated just below the PTBP1 complex (Fig. 1D, lane 2, asterisk). We also used an unrelated oligonucleotide with the sequence identical to the first 20 nucleotides of U7 snRNA (5’ U7), containing the CUCUUU SBP site (Fig. 1A). This oligonucleotide only partially inhibited the formation of the PTBP1 complex (Fig. 1D, lane 3), suggesting that it has a lower affinity for PTBP1. Interestingly, the two oligonucleotides while preventing or limiting the interaction of U7/WT RNA with PTBP1, stimulated binding of this RNA to Gemin5 (Fig. 1D, lanes 3-6, arrow). This result suggests that the competition with PTBP1 may be one of the reasons for the inability of Gemin5 to bind to the Sm site in U7 snRNA (see below).

### PTBP1 binds at least two independent sites in U7 snRNA

To analyze binding of PTBP1 to U7 using a more defined system, the L and S isoforms of PTBP1 were separately synthesized *in vitro* in the presence ^35^S methionine using the TnT system (Promega), and the labeled proteins were resolved by SDS/PAGE and detected by autoradiography (Fig. 2A, top panel, lanes 2 and 3). As determined by Western blotting, the L and S isoforms co-migrated with endogenous isoforms from HeLa cells (Fig. 2A, bottom panel, compare lanes 1 and 3), supporting the notion that the longer form contains additional 24 amino acids between RRM2 and RRM3. A control sample (no exogenous cDNA used) shifted a small fraction of U7/WT probe, (Fig. 2B, lanes 3 and 6), likely due to the presence of endogenous PTBP1 and/or proteins with similar binding properties in the lysate used for *in vitro* translation. Each of the two ^35^S-labeled PTBP1 splice variants formed a much more pronounced complex co-migrating with the cytoplasmic complex (Fig. 2B, lanes 4 and 7). Importantly, the TnT-generated PTBP1 failed to interact with U7/SPL (Fig. 1B, lane 10), hence mimicking the behavior of endogenous PTBP1 from HeLa cytoplasmic extracts.

**Fig. 2.**
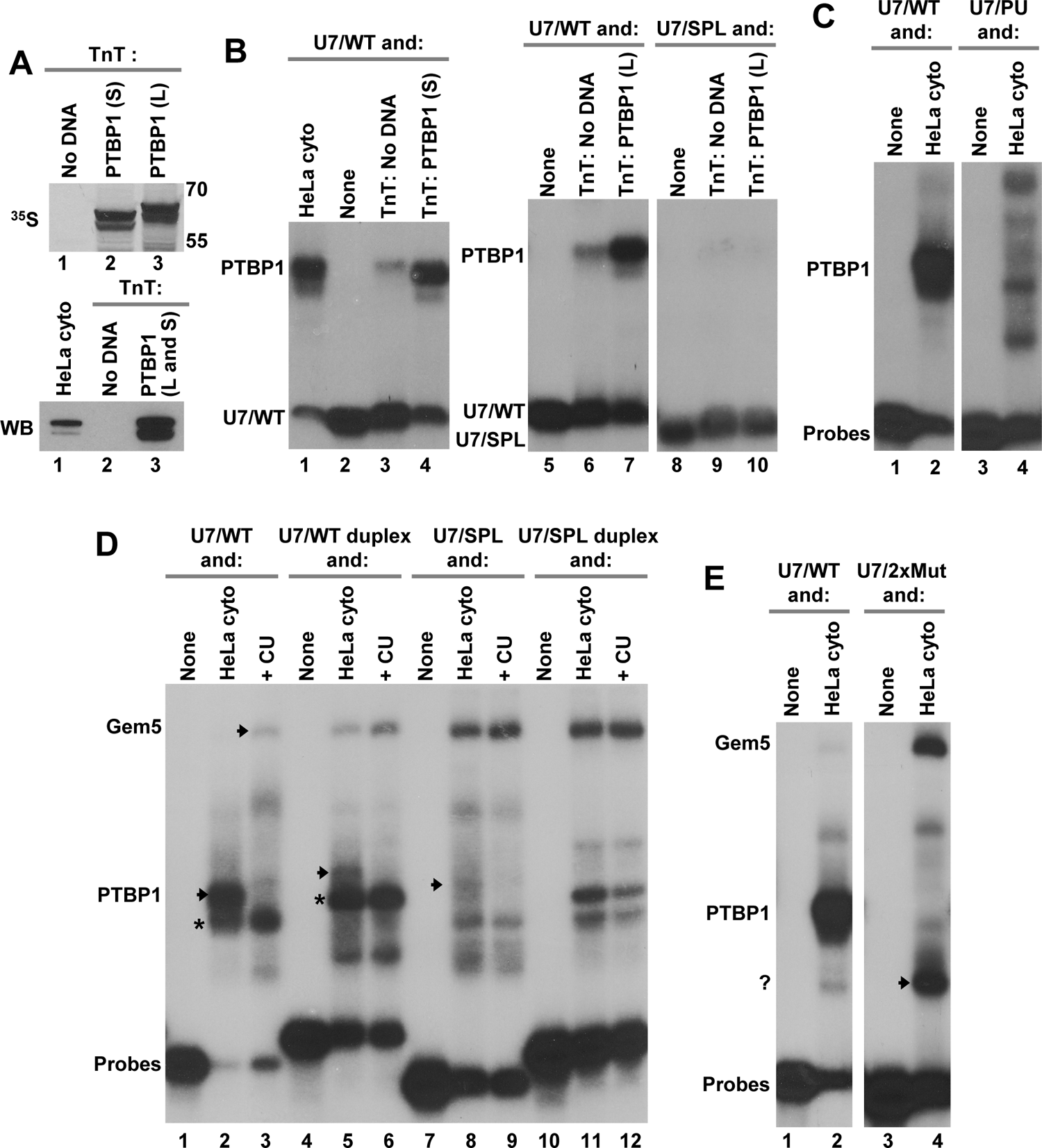
PTBP1 directly binds U7 snRNA in a manner that depends on the SBP and Sm sites. **A.** The two isoforms of PTBP1, short (S) and long (L), were separately expressed in rabbit reticulocyte lysate in the presence of ^35^S using the TnT system, separated by PAGE and detected by autoradiography (top) or Western blotting next to a small aliquot of a HeLa cytoplasmic extracts (lane 1, bottom). *In vitro* translation in the absence of cDNA is shown in lanes 1 (top) and 2 (bottom). In lane 3 of the bottom panel, the two *in vitro* generated isoforms were mixed. **B and C.** EMSA with indicated probes and cytoplasmic extracts or TnT-expressed PTBP1 (short and long isoform), as described in the legend for Fig.1. **D.** EMSA with a HeLa cytoplasmic extract and U7/WT or U7/SPL probes alone or pre-annealed to an oligonucleotide complementary to the first 18 nucleotides of U7 snRNA (duplexes). The arrows indicate positions of the PTB1- and Gemin5-containing complexes, and asterisks in lanes 2 and 5 indicate a complex that migrates just below the PTBP1-complex. In lanes 3, 6, 9 and 12, the extract contains excess of the CU oligonucleotide to sequester endogenous PTBP1. The position of PTBP1- and Gemin5-containing complexes is indicated with the arrows. **E.** EMSA with a HeLa cytoplasmic extract and indicated probes. The question mark in panel E indicates a complex that likely contains hnRNP A1.

In addition to PTBP1, we also tested *in vitro* generated PTBP2. This neuronal-specific paralog shares 75% sequence identity with PTBP1 and displays the same binding properties, forming a complex with U7/WT but not with U7/SPL (not shown). Collectively, these results strongly argue that PTBP1 and PTBP2 specifically recognize the unique Sm site in U7 snRNA and have a much weaker affinity to U7 snRNA containing a spliceosomal-type Sm site.

Upstream of the Sm site, all known U7 snRNAs contain a CUCU sequence, a likely binding site for PTBP. This sequence is part of the larger CUCUUU substrate recognition site that base pairs with the purine rich HDE in histone pre-mRNAs and therefore plays a key role in recruiting substrates for U7 snRNP (7,8). We changed the CUCU to complementary purines, GAGA, hence creating U7/PU RNA and analyzed its ability to bind PTBP1 using EMSA. Also this mutant RNA failed to form a complex with both the cytoplasmic (Fig. 2C, lane 4) and *in vitro* generated PTBP1 (not shown). U7/WT RNA tested in parallel readily formed a complex with PTBP1 from the same extract (Fig. 2C, lane 2). Thus, PTBP1 simultaneously recognizes two functionally essential and CU-containing sequence elements in U7 snRNA, the SBP and Sm sites. The 4-nucleotide substitution that converted U7/WT into U7/PU RNA failed to result in detectable Gemin5 binding, likely due to incomplete inhibition of PTBP1 binding A (not shown).

We next investigated the effect of blocking the 5’ region of U7 snRNA (nucleotides 1-18) by an antisense oligonucleotide. Clearly, forming a duplex between these two RNAs eliminates the CUCU single stranded region that binds PTBP1, but could also change the overall folding of U7 snRNA, hence affecting its interaction with other proteins. Consistent with the data shown above, U7/WT probe interacted with PTBP1 from a HeLa cytoplasmic extract, generating a stable complex migrating near the middle of the gel (Fig. 2D, lane 2). In the presence of molar excess of the CU oligonucleotide, the PTBP1 complex was eliminated, and the complex containing Gemin5 became more apparent (Fig. 2D, lane 3, arrow), in agreement with the data shown in Fig. 1D, lane 3, suggesting that the inability of Gemin5 to bind U7/WT at least in part results from the competition with PTBP1. The presence of the CU competitor also demonstrated that what appeared to be a single PTBP1 complex, consists in fact of two closely migrating complexes, with only the top complex containing PTBP1 and the bottom one containing a different protein insensitive to the CU oligonucleotide (asterisk). As expected, given the importance of the CUCU element, U7 snRNA duplexed with the anti-sense oligo, only weakly interacted with PTBP1 (Fig. 2D, lane 5, arrow). Note that the duplex formation shifts each probe and its complexes to higher positions on the gel. Again, the Gemin5-containing complex was formed, and the intensity of this complex increased in the presence of the CU competitor (Fig. 2D, lane 6). U7/SPL RNA retained some ability to bind cytoplasmic PTBP1, as expected given the presence of the second binding site in this mutant (Fig. 2D, lane 8, arrow). Importantly, this residual binding was abolished in the presence of the CU oligonucleotide, resulting at the same time in a detectable increase in the intensity of the Gemin5 complex (Fig. 2D, lane 9). One interpretation of these results is that while the CU oligonucleotide prevents any interaction between U7/WT and PTBP1, the anti-sense oligonucleotide affects folding of the RNA, with the two effects independently contributing to converting U7/WT from a preferable target for PTBP1 to a target that detectably interacts with Gemin5.

In addition to partially explaining the inability of Gemin5 to interact with U7/WT RNA, the above experiment confirmed that PTBP1 recognizes at least two independent sequence elements in U7 snRNA: substrate base-pairing (SBP) site and Sm site. We next designed an RNA in which these two sites were simultaneously mutated and tested the resultant U7/2xMut RNA in a mobility shift assay using a HeLa cytoplasmic extract. As expected, most of the U7/WT probe was shifted in the native gel to a higher position due to binding cytoplasmic PTBP1, whereas U7/2xMut probe tightly interacted with Gemin5, failing to form a complex with PTBP1 (Fig. 2E, lanes 2 and 4). In addition, a pronounced complex was detected close the bottom of the gel (question mark). This complex is likely formed by hnRNP A1 and its paralogues (see below). These proteins, as revealed by affinity purification experiments, readily interact with mutant RNAs unable to bind PTBP1 (see for example Fig. 1B, lanes 2 and 3).

### Recombinant PTBP1 binds U7 snRNA as a dimer

We bacterially expressed PTBP1 (longer form) N-terminally tagged with either five histidines (5xH) or glutathione S-transferase (GST). The two recombinant proteins were tested for their ability to bind U7/WT probe in EMSA. Surprisingly, each protein formed with U7/WT probe a complex migrated significantly higher than the cytoplasmic complex containing endogenous PTBP1 (Fig. 3A, lanes 2-4). One possibility was that the two recombinant proteins form dimers due to their large molar excess over the probe. This possibility was supported by the presence of a small amount of a 5xH-PTBP1 complex that migrated with as similar mobility as the cytoplasmic complex (Fig. 3A, lane 3, arrow).

**Fig. 3.**
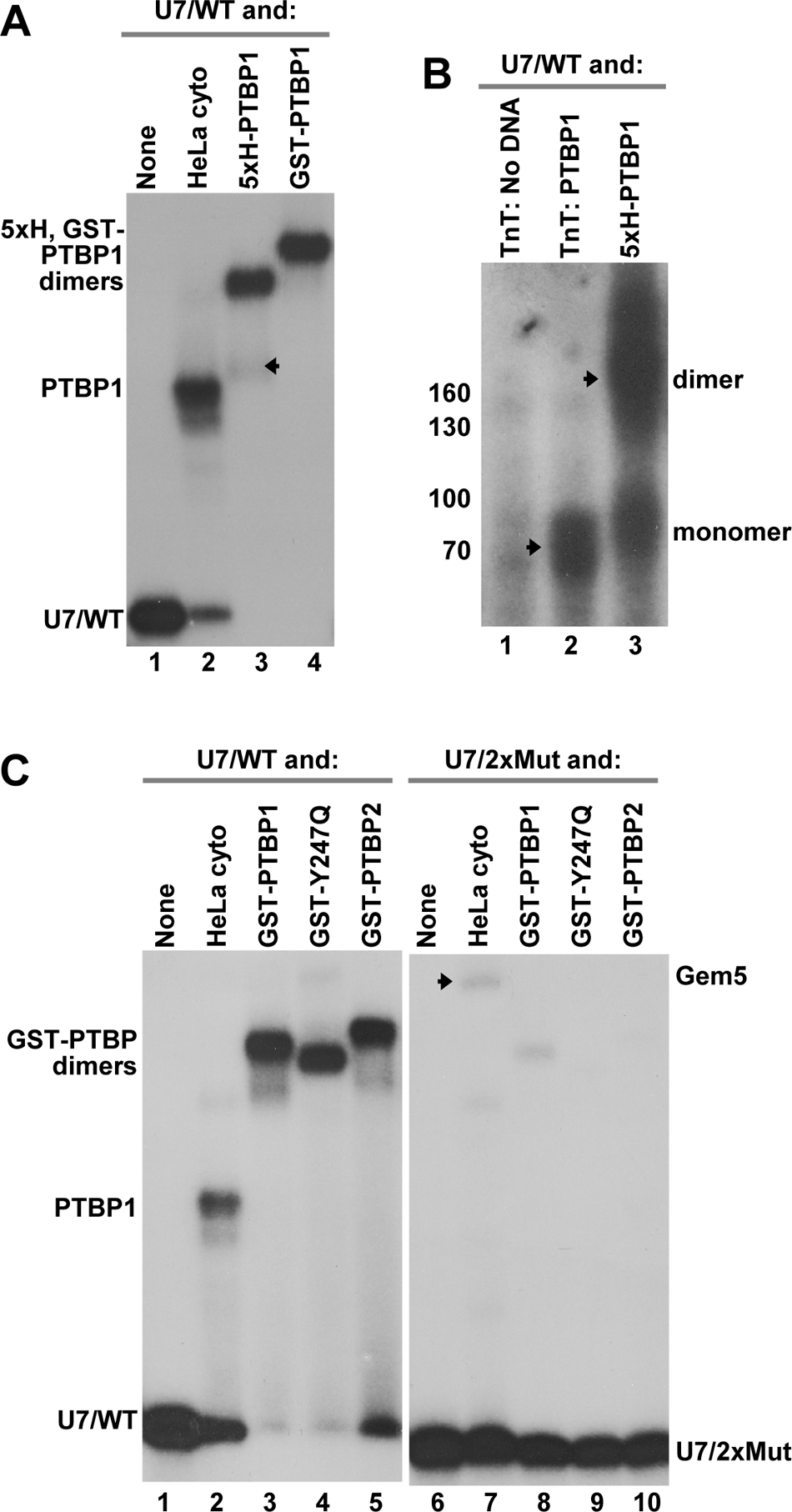
Binding of recombinant PTBP1 to U7 snRNA. **A.** EMSA with U7/WT probe and a Hela cytoplasm or indicated recombinant proteins N-terminally tagged with 5xH(istidine) or GST. The position of PTBP1 monomer in complex with U7/WT is indicated with an arrow. **B.** TnT-expressed PTBP1 (lane 2) or 5xH-tagged recombinant PTBP1 (lane 3) were UV irradiated and separated by PAGE. Proteins covalently attached to radioactively labeled RNA were detected by autoradiography. In lane 1, UV-crosslinking was carried out with a control TnT lysate lacking PTBP1 cDNA. The position of the PTBP1 monomer and dimer cross-linked to U7/WT probe is indicated. **C.** EMSA with U7/WT or U7/2xMut probes and a Hela cytoplasm or indicated recombinant proteins. The arrow in lane 7 indicates the position of Gemin5.

PTBP1 was previously shown to dimerize under nonreducing conditions via cysteine 23 located in the N-terminal region preceding RRM1 (80,81). The presence of a high concentration of DTT did not significantly reduce the amount of the large PTBP1 complex (data not shown), suggesting that the two recombinant proteins may multimerize upon binding U7 snRNA (80,81), with the large molar excess of each protein over the probe enhancing PTBP1 dimerization. Our attempts to lower the amount of recombinant PTBP1 resulted in its precipitation.

Whether recombinant proteins form dimers was next investigated by UV-cross-linking. In these experiments, U7/WT radioactive probe was mixed with a control lysate (no cDNA used for *in* vitro translation), *in vitro* expressed PTBP1 or recombinant 5xH-tagged PTBP1, and the samples were UV irradiated. Proteins covalently attached to the RNA were separated by SDS/PAGE without a prior treatment with RNases to prevent the removal of the 5’ ^32^P, and radioactively labeled proteins were detected by autoradiography. Note that the RNA moiety (20 kDa) increases the molecular size of monomeric PTBP1 from ∼60 kDa to ∼80 kDa. While UV-cross linking of WT/U7 probe with the control lysate failed to yield major protein bands with the radioactive label (Fig. 4B, lane 1), a single diffused band of ∼80 kDa was detected in the presence of *in vitro* expressed PTBP1. Recombinant 5xH-PTBP1 yielded both the 80 kDa cross-link and a larger species migrating at ∼160 kDa size marker, indicative of cross-linking two protein molecules to the same RNA molecule.

**Fig. 4.**
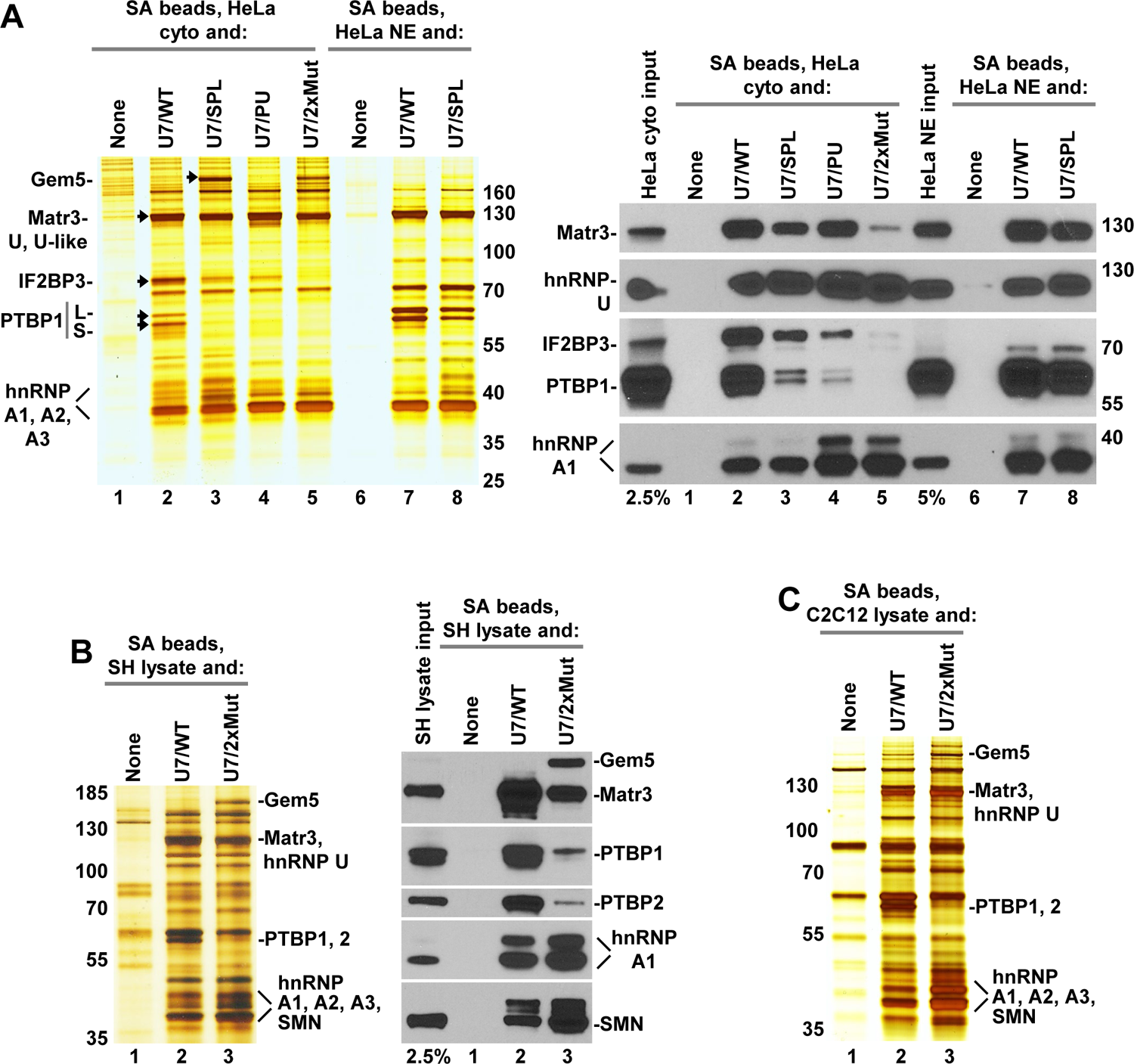
The ubiquitous nature of the PTBP1/U7 snRNA interaction. **A.** Affinity purification of proteins from a HeLa cytoplasmic extract (lanes 1-5) and a nuclear extract (NE) (lanes 6-8). Proteins bound to U7/WT RNA or its mutants, as indicated, were immobilized on SA beads, separated in a 4-12% SDS/polyacrylamide gel and analyzed by silver staining (left panel) or Western blotting (right). Lane 1 contains proteins that bind to SA beads in the absence of RNA. In the right panel, the input lanes contain ∼2.5% and ∼5% of the cytoplasmic and nuclear extracts used in the binding experiments, respectively. **B and C.** Affinity purification of proteins from human SH-SY5Y neuroblastoma cells (panel B) and mouse C2C12 skeletal muscle cells (panel C). Proteins bound to U7/WT or U7/2xMut RNAs were visualized by silver staining and Western blotting (B, left and right, respectively) or only silver staining (C).

When tested by EMSA, 5xH-PTBP1 and GST-PTBP1 also bound mutant probes, U7/SPL and U7/PU (not shown). This clearly resulted from using an excess of recombinant proteins and the fact that each mutant RNA contains more than one PTBP1 recognition site. We bacterially expressed two additional GST-tagged recombinant proteins: PTBP2 and a mutant version of PTBP1 containing a Y247Q substitution known to abolish interaction of PTBP1 with Matrin3 (59) (see below). The Y247Q PTBP1 also lacked the entire unstructured N-terminus, including cysteine 23 that mediates intermolecular disulfide bond formation and PTBP1 dimerization in nondenaturing environments (80).

In agreement with the data shown in Fig. 3A, GST-tagged PTBP1, Y247Q PTBP1 and PTBP2 shifted U7/WT probe to a position in the gel consistent with each recombinant protein forming a dimer with the RNA (Fig. 3C, lanes 3-5). Note that Y247Q PTBP1 is unable to form a disulfide bond due to the lack of cysteine 23. Thus, dimerization of this mutant is likely triggered by RNA binding and high protein concentration used in the assay. Importantly, none of the three recombinant proteins bound U7/2xMut probe in which the two CU-containing sites, SBP and Sm, were mutated (Fig. 4C, lanes 8-10). We conclude that bacterially expressed PTBP1 and PTBP2 bind U7 snRNA as dimers and simultaneously recognize two sequence elements that are essential for the function of U7 snRNP.

### The ubiquitous nature of the PTBP1/U7 snRNA interaction

Having established that PTBP1 is virtually unable to interact with U7/2xMut RNA even in the presence of high protein concentrations, we used this RNA as a negative control in several affinity purification experiments with various mammalian extracts. Compared to the initial experiments, we increased the amount of RNA 10-fold (7.5 µg of RNA for 750 µl of extract) to purify proteins with both high and low affinity for U7 snRNA. Consistent with the initial experiments, silver staining (Fig. 4A, left) and Western blotting (Fig. 4A, right) identified only two proteins in a HeLa cytoplasm that were affected by combining mutations within SBP and Sm sites: PTBP1 and IGF2BP3 (Fig. 4A, both panels, lane 2). Binding of these proteins to U7 snRNA was significantly reduced by individually mutating each of the two binding sites (Fig. 4A, both panels, lanes 3 and 4), and was virtually eliminated by mutating the two sites simultaneously (Fig. 4A, both panels, lane 5). The two RNAs with a spliceosomal-type Sm site, U7/SPL and U7/2xMut, readily interacted with cytoplasmic Gemin5, confirming the data from EMSA. Other HeLa cytoplasmic proteins, including hnRNP U, U-like1 and hnRNP A1, bound each of the three mutant RNAs with the same or higher efficiency than did U7/WT. As determined by Western blotting, binding of Matrin3, which is recruited to U7 snRNA via PTBP1, was almost entirely abolished by mutating the two sites simultaneously (Fig. 4A, right, lane 5). The effect of each individual mutation was masked by the strong signal produced by the anti-Matrin3 antibody (Fig. 4, right, lanes 3 and 4).

We also used U7/WT and U7/SPL RNAs to test whether the same binding pattern will be observed with a nuclear extract prepared from the same batch of HeLa cells. In contrast to the cytoplasmic extract, the nuclear extract contained a significantly lower concentration of Gemin5 (not shown), and IGF2BP3 was undetectable (Fig. 4, right, NE input), consistent with the cytoplasmic localization of this protein (82–84). Based on silver staining, U7/SPL RNA was less efficient in binding PTBP1 from nuclear extract (Fig. 4, left, compare lanes 7 and 8), although Western blotting failed to detect this difference due to the strong signal produced by the antibody (Fig. 4A, right, lanes 7 and 8). We conclude that PTBP1 and IGF2BP3 are the only HeLa proteins that in our affinity purification assays display specific interaction with wild type U7 snRNA but not with U7 snRNA mutated within the SBP and Sm sites.

We next used human SH-SY5Y neuroblastoma (85) and mouse C2C12 skeletal muscle cells to determine whether binding of PTBP1 and IGF2BP3 is conserved across different mammalian species and tissues. We choose to prepare whole cell lysates from these cells to simultaneously look for U7 snRNA binders in both the cytoplasmic and nuclear fractions and used two RNAs with drastically different binding properties: U7/WT and U7/2xMut. The two cell lines, despite representing two different organisms and tissue types, yielded essentially the same results (Fig. 4B and 4C). Of the complex pattern of band visualized by silver staining, only one band was specific for each RNA: PTBP1 for U7/WT RNA, and Gemin5 for U7/2xMut. The material bound to U7/WT RNA also contained significant amounts of PTBP2. We conclude that PTBP1/2 and Gemin5 are the only proteins that under our experimental condition specifically interact with U7/WT and U7/2xMut, respectively.

Surprisingly, Western analysis of the material purified from the two cell lysates by U7/WT revealed the presence of the SMN protein (Fig. 4B, right, lanes 2 and 3). As expected, a significantly larger amount of this protein was detected in the material purified via U7/2xMut, as this RNA interacts with Gemin5, which in turn recruits some other subunits of the SMN complex, including the SMN protein.

### Binding of hnRNP A1 to U7 snRNA

One of the RNA binding proteins affinity-purified by both the wild type U7 snRNA (U7/WT) and its three mutants, U7/SPL, U7/PU and U7/2xMut, was hnRNP A1 and the two paralogues, hnRNP A2 and A3. In fact, the interaction of these proteins with U7 snRNA was clearly improved by mutations that disrupted binding of PTBP1, suggesting that PTBP1 and hnRNP A1 compete for the same or overlapping binding sites. Previous structural and biochemical studies demonstrated that hnRNP A1 recognizes UAG trinucleotides (55), and two of them are present in U7 snRNA, one separating the SBP and Sm sites and the other being located at the 3’ end of the Sm site (Fig. 1A). This latter UAG motif is the hallmark of the Sm site and overlaps with the preceding motif that interacts with PTBP1. The two UAG trinucleotides are universally conserved in vertebrate U7 snRNA, but surprisingly are absent from the *Drosophila* U7 snRNA (Fig. 5A).

**Fig. 5.**
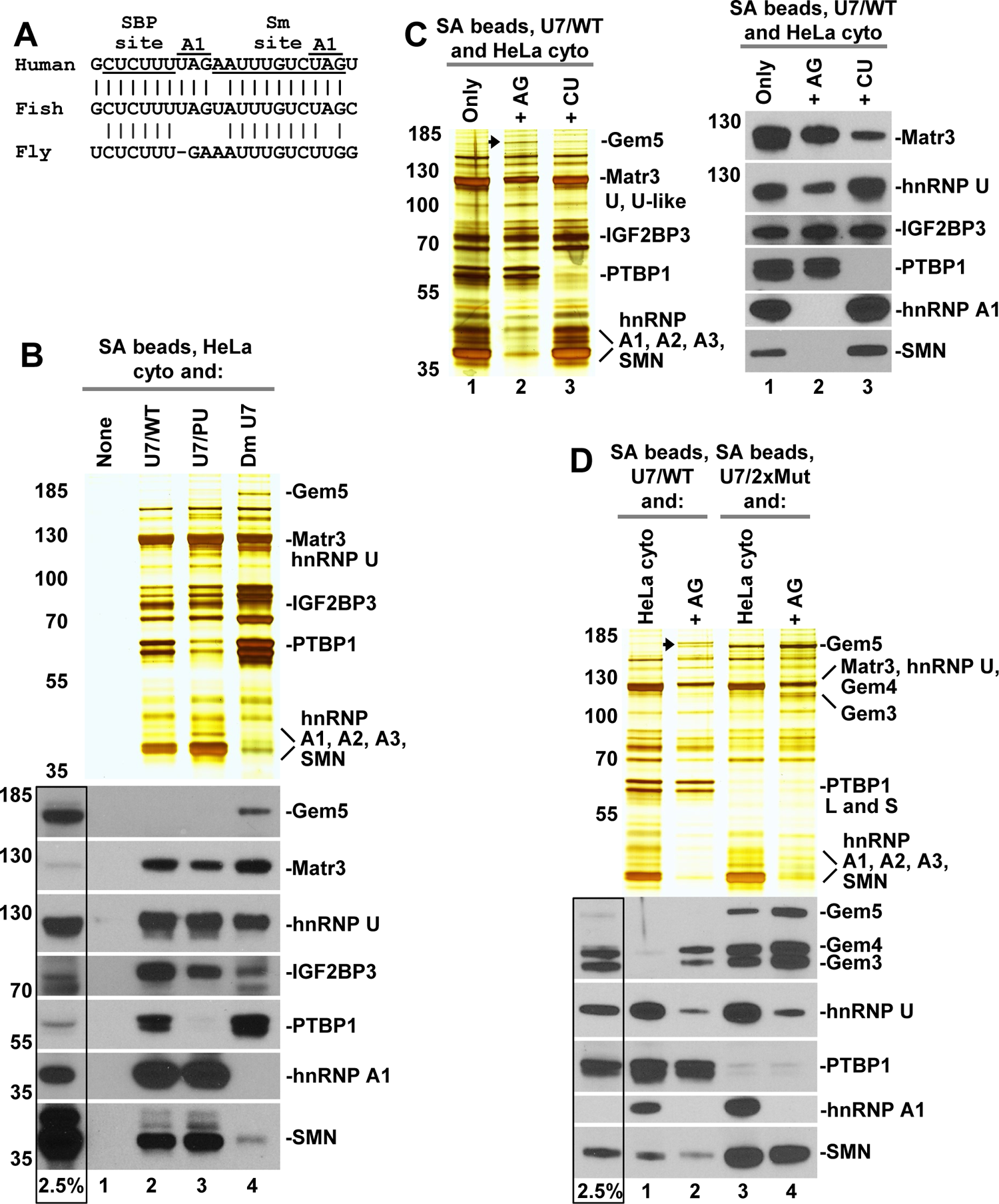
Binding of hnRNP A1 to U7 snRNA. **A.** Sequence alignment of the SBP and Sm sites in U7 snRNA from *H. sapiens*, *D. rerio* and *D. melanogaster*. The two sites are underlined and the two UAG motifs predicted to bind hnRNP A1 are indicated. **B.** Proteins from a HeLa cytoplasmic extract bound to indicated RNAs were immobilized on SA beads, separated in a 4-12% SDS/polyacrylamide gel and analyzed by silver staining (top) or Western blotting (bottom). Lane 1 contains cytoplasmic proteins immobilized on SA beads in the absence of RNA. In the bottom panel, the input lane (boxed) contains ∼2.5% of the cytoplasmic extract used in the binding experiment. **C.** Affinity purification of proteins bound to U7/WT RNA from a HeLa cytoplasmic extract (lane 1) or from the same extract pre-incubated with excess of the AG (lane 2) or CU oligonucleotide competitors (lane 3). Proteins immobilized on SA beads were detected by silver staining (left panel) or Western blotting (right panel). **D.** Affinity purification of proteins bound to either U7/WT or U7/2xMut RNAs, as indicated, in an untreated HeLa cytoplasmic extract (lanes 1 and 3) or in the same extract pre-incubated with excess of the AG oligonucleotide (lanes 2 and 4). Proteins immobilized on SA beads were detected by silver staining (top) or Western blotting (bottom). The arrow indicates the band that corresponds to Gemin5. In the bottom panel, the input lane (boxed) contains ∼2.5% of the cytoplasmic extract used in the binding experiment.

To facilitate mapping the regions of U7 snRNA that are essential for binding hnRNP A1, we compared proteins of HeLa cytoplasmic extract that interact with U7/WT, U7/PU and *Drosophila* (Dm U7) U7 snRNAs. Human and *Drosophila* U7 snRNAs are partially conserved only within a short region encompassing the SBP and Sm sites, with the two UAG motifs in vertebrate U7 snRNA being replaced in *Drosophila* U7 snRNA with UGA and UUG (Fig 5A). Beyond the SBP and Sm sites, human and *Drosophila* U7 snRNAs have no recognizable homology. Consistent with the data shown above, U7/WT RNA was more efficient than U7/PU RNA in binding PTBP1 (and hence its binding partner Matrin3) and IGF2BP3, whereas U7/PU RNA, as judged by silver staining, bound more hnRNP A1. *Drosophila* U7 snRNA, despite its vastly different sequence exceeded U7/WT in its ability to ability to interact with PTBP1 but failed to interact with IGF2BP3 and hnRNP A1. Strikingly, Dm U7 also interacted with Gemin5 (Fig 5B, lane 4), strongly contrasting with U7 snRNA. Intriguingly, while failing to bind Gemin5, U7/WT and U7/PU RNAs were both associated with SMN, with the U7/PU-bound material containing more of this subunits, in a good correlation with the increased amount of hnRNP A1 (Fig 5B, compare lanes 2 and 3). On the other hand, the material bound to Dm U7 snRNA contained a barely detectable amount of SMN, correlating with the amount of Gemin5 (Fig 5B, compare lanes 2 and 3). These results suggest that depending on the sequence of the Sm site, the SMN protein can be recruited by two independent mechanisms, via Gemin5 (spliceosomal Sm site) or hnRNP A1 (U7-specific Sm site).

We reasoned that the inability of Gemin5 to interact with human U7 snRNA might result from a tight interaction of hnRNP A1 to this RNA, resembling the same competition mechanism observed for PTBP1. Note that the two UAG high affinity targets for hnRNP A1 in U7 snRNA flank the key nucleotides in the Sm site known to bind Gemin5 (24,86–89). The lack of this competition could therefore allow Gemin5 to find its binding site in *Drosophila* U7 snRNA. To test this possibility, we used excess (15 µg) of the AG oligonucleotide containing high affinity binding sites for hnRNP A1(90) to sequester this protein in 750 µl of a HeLa cytoplasmic extract. In parallel, the same amount of the extract was incubated with the CU oligonucleotide to sequester PTBP1. These two extracts and an untreated extract were next incubated with U7/WT RNA and bound proteins were immobilized on streptavidin beads and analyzed by silver staining and Western blotting. As expected, U7/WT RNA bound PTBP1 in the control extract but not in the extract containing excess of the CU oligonucleotide (Fig. 5C, left and right panels, lanes 1 and 3, respectively). In the presence of this oligonucleotide, binding of hnRNP A1 and IGF2BP3 was virtually unaffected. The UAG oligonucleotide acted in the opposite manner, preventing the interaction of U7/WT RNA with hnRNP A1 but having no effect on the interaction with PTBP1 (Fig. 5C, left and right panels, lane 2). In addition, this oligonucleotide significantly inhibited binding of hnRNP U to this RNA. Importantly, the UAG oligonucleotide eliminated SMN from the material bound to U7/WT RNA (Fig. 5C, right panel, lane 2), suggesting that this subunit of the SMN complex is indirectly recruited to U7 snRNA via hnRNP A1.

The UAG oligonucleotide, besides preventing hnRNP A1 from interacting with U7/WT, resulted in binding of a protein migrating at ∼170 kDa (Fig. 5C, left panel, lane 2, arrow), most likely Gemin5, although Western blotting failed to confirm this identity due to the small amount of the protein (not shown).

We carried out a similar experiment with a HeLa cytoplasmic extract and the AG oligonucleotide, additionally including WT/2xMut RNA, which binds Gemin5 due to the presence of a spliceosomal-type Sm site. In the material bound to WT/U7, hnRNP A1 was readily detectable and Gemin5 was absent (Fig. 5D, lane 1). When binding was carried out with HeLa extract pre-incubated with the AG competitor, hnRNP A1 was missing, and a protein migrating with the mobility of Gemin5 was detected by silver staining, mirroring the results shown in Fig. 5B. Although the anti-Gemin5 antibody again failed to confirm this identity, antibodies against Gemin3 and Gemin4 clearly demonstrated that these two subunits of the SMN complex were enriched in the presence of the AG oligonucleotide. The same effect was observed for WT/2xMut RNA. While this RNA due to the presence of a spliceosomal-type Sm site binds Gemin5 on its own, this component of the SMN complex and its two binding partners, Gemin3 and Gemin4, were clearly enriched in the presence of the AUG oligonucleotide (Fig. 5D, compare lanes 3 and 4). Collectively, these results suggest that binding of Gemin5 to the Sm site U7 snRNA is at least in part limited by competition with hnRNP A1.

To provide more direct evidence that binding of hnRNP A1 to human U7 snRNA depends on the two UAG motifs located in the proximity of the Sm site, we switched the position of purines in these trinucleotides to UGA (Fig. 1A). The resultant mutant U7 snRNA, U7/2xGA, when incubated with a HeLa cytoplasmic extract only weakly bound hnRNP A1 (Fig. 6A, compare lanes 2 and 3). The same effect was observed for SMN, supporting the notion that this protein is indirectly recruited to U7 snRNA by hnRNP A1 bound to the two UAG motifs located in the vicinity of the Sm site.

**Fig. 6.**
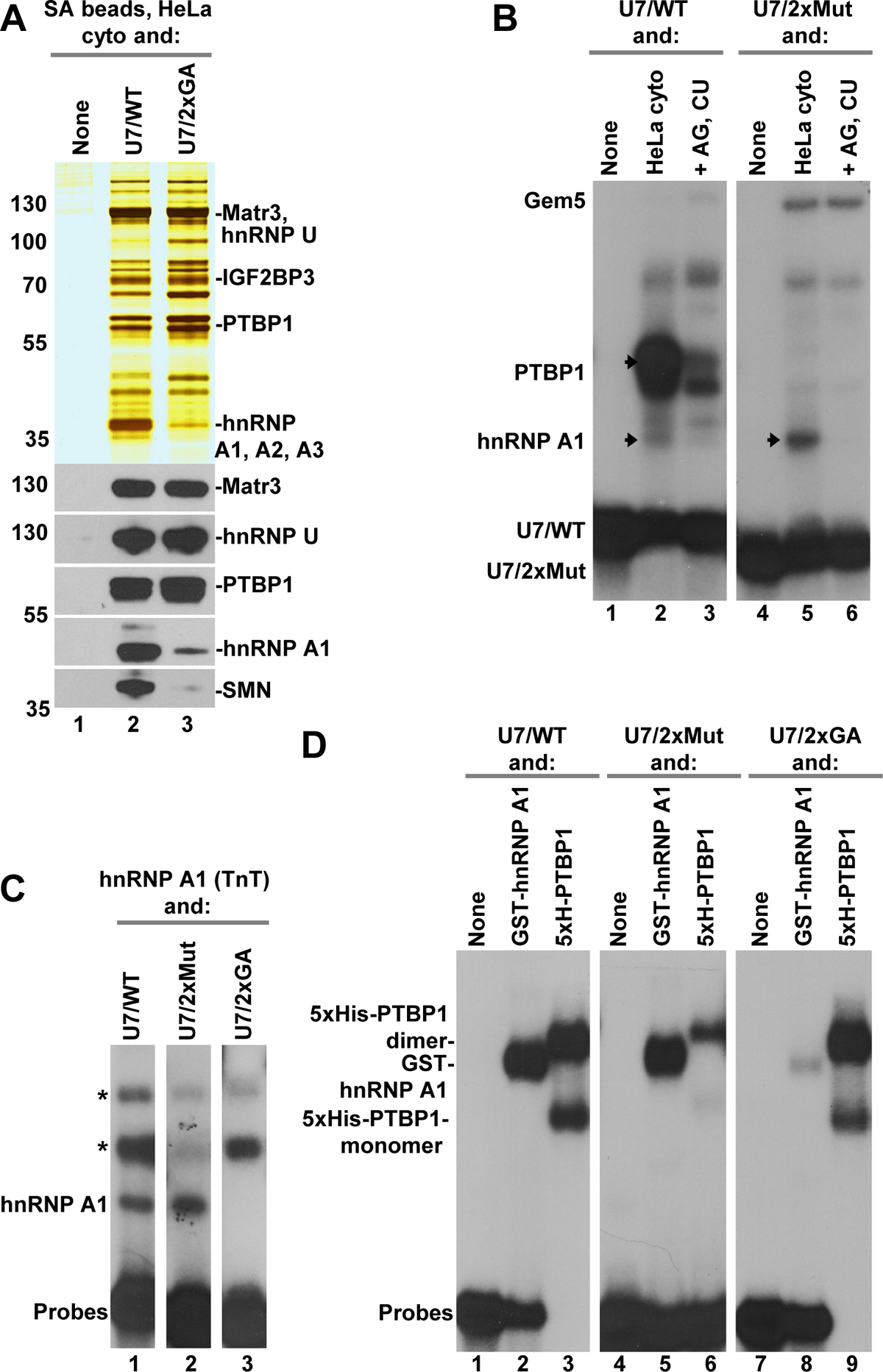
hnRNP A1 requires two UAG motifs flanking the Sm site in U7 snRNA. **A.** Affinity purification of proteins from a HeLa cytoplasmic extract bound to U7/WT or U7/2xGA RNAs. Proteins immobilized on SA beads were detected by silver staining (top) or Western blotting (bottom). Lane 1 contains cytoplasmic proteins immobilized on SA beads in the absence of RNA. **B.** EMSA with U7/WT or U7/2xMut probes incubated in a Hela cytoplasm or the same extract pre-incubated with excess of AG and CU oligonucleotides. Arrows in lanes 2 and 5 indicated protein complexes containing hnRNP A1. **C.** EMSA with indicated probes and TnT-expressed hnRNP A1. The asterisks indicate complexes containing proteins from the TnT lysate, with the bottom complex likely containing endogenous PTBP1. **D.** EMSA with indicated probes and recombinant GST-hnRNP A1 or 5xH-PTBP1.

We next investigated whether the interaction between hnRNP A1 and U7 snRNA can be detected by EMSA. As shown above, U7/WT probe mixed with a HeLa cytoplasmic extract yields a major complex containing PTBP1, and a few minor complexes of unknown origin. To determine whether any of these complexes contains hnRNP A1, we pre-incubated the cytoplasmic extract with a 500-fold molar excess of two oligonucleotides, CU and AG, to simultaneously sequester PTBP1 and hnRNP A1, respectively. Only two complexes were eliminated (Fig. 6B, lane 3): the major complex containing PTBP1 and a barely detectable complex migrating below, likely containing hnRNP A1 (Fig. 6B, compare lanes 2 and 3, arrow). This conclusion was confirmed by using U7/2xMut RNA, which is unable to interact with PTBP1 but forms a much more pronounced lower complex (Fig. 6B, lane 5, arrow, see also Fig. 2E, lane 4, arrow). This lower complex was eliminated in the presence of oligonucleotides, confirming that it contains hnRNP A1. The electrophoretic mobility of this complex suggests that endogenous hnRNP A1 interacts with U7 snRNA as a monomer.

We also expressed hnRNP A1 by *in vitro* translated and tested ^35^S-labeled protein by EMSA. Of three probes used, U7/WT and U7/2xMut (Fig. 6C, lanes 1 and 2), but not U7/2xGA (Fig. 6C, lane 3), formed a complex migrating near the bottom of the gel. Finally, we bacterially expressed GST-tagged hnRNP A1 and tested its ability to bind various U7 RNA probes by EMSA, using recombinant 5xH-tagged PTBP1 as a specificity control. U7/WT interacted with both recombinant proteins, as expected (Fig. 6D, lanes 2 and 3). Note that 5xH-PTBP1 forms with U7/WT RNA both a monomer and a dimer. Based on relative mobilities, we tentatively conclude that GST-tagged hnRNP A1 forms a dimer, likely resulting from the high concentration of the recombinant protein. Of the two mutant probes used, U7/2xMut was selectively impaired in binding PTBP1 (Fig. 6D, lane 6), as expected, whereas U7/2xGA was unable to bind GST-hnRNP A1 but was unaffected in forming a complex with PTBP1 (Fig. 6D, lanes 8 and 9). We conclude that hnRNP A1 directly interacts with human U7 snRNA, and for binding requires two highly conserved UAG motifs flanking the Sm site.

### Recombinant PTBP1 and endogenous hnRNP A1 can form a ternary complex with U7 snRNA

We took advantage of the availability of recombinant GST-tagged PTBP1 to identify its binding partners in HeLa cell extracts. GST-PTBP1 either alone or in a complex with U7 snRNA was incubated with a HeLa nuclear extract and the bound proteins were immobilized on glutathione (GSH) beads and analyzed by silver staining (Fig. 7A). Only two major background proteins were detected by silver staining when GSH beads were incubated with the nuclear extract alone (Fig. 7A, top, lane 1). Strikingly, the GST-PTBP1 in a complex with U7/WT RNA (Fig. 7A, top, lane 3), but not GST-PTBP1 itself (Fig. 7A, top, lane 2), bound to a group of nuclear proteins migrating around 38 kDa. Analysis by mass spectrometry determined that this band contains hnRNP A1 and its closely related paralogues, hnRNP A2 and A3. The identity of hnRNP A1 was confirmed by Western blotting (not shown). The ternary complex did not contain hnRNP U1 (not shown), a protein that binds both wild type U7 snRNA and its mutants, suggesting that the association of hnRNP A1 is specific. Importantly, Western blotting revealed the material purified by the GST-PTBP1/RNA complex, but not by the protein itself, contains detectable amounts of SMN (Fig. 7A, bottom, lane 3), supporting the notion that this subunit of the SMN complex interacts with hnRNP A1 or its paralogues.

**Fig. 7.**
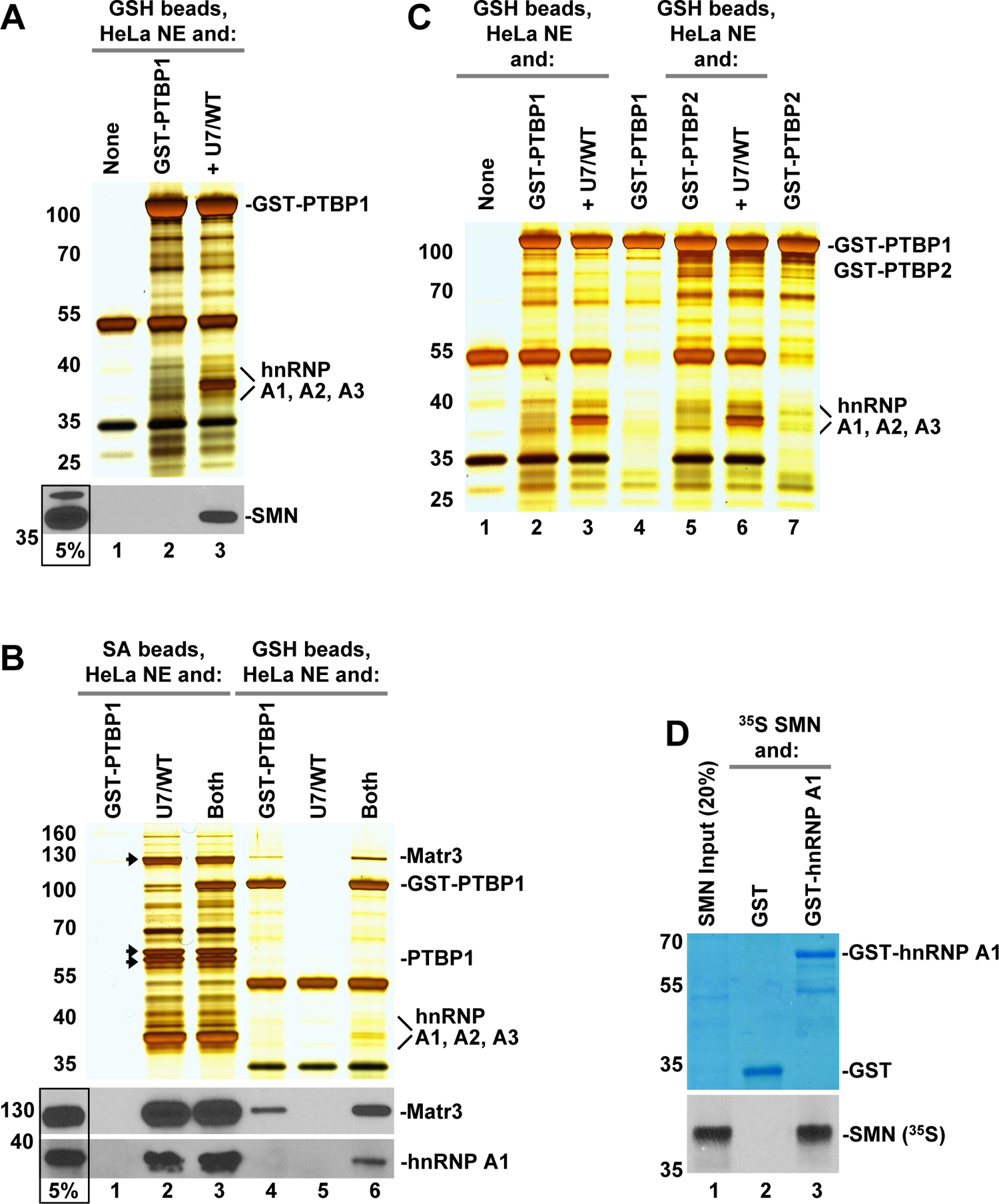
PTBP1 and hnRNP A1 form a ternary complex with U7 snRNA. **A.** Recombinant GST-tagged PTBP1 alone or pre-bound to U7 snRNA were incubated with a HeLa nuclear extract (NE) and proteins bound to GST-PTBP1 were immobilized on glutathione (GSH) beads and analyzed by silver staining (top). The presence of SMN was monitored by Western blotting (bottom). **B.** Recombinant GST-tagged PTBP1 alone, U7/WT RNA alone, or a complex of both of them were incubated with a HeLa nuclear extract (NE). The samples were divided into two haves and each half incubated with streptavidin (SA) or glutathione (GSH) beads and immobilized proteins analyzed by silver staining (top). The presence of hnRNP A1 and Matrin3 was monitored by Western blotting (bottom). Silver-stained bands corresponding to Matrin3 and the two forms of PTBP1 are indicated with arrows. **C.** As in panel A, with the exception that the experiment additionally included GST-tagged PTBP2, and the two recombinant proteins were additionally immobilized on GSH beads in the presence of buffer (lanes 4 and 7). **D.** *In vitro* generated ^35^S-labeled SMN was incubated with GST or GST-tagged hnRNP A1 and the bound proteins were immobilized on GSH beads. GST proteins were visualized by staining with Coomassie Blue (top), and ^35^S-labeled SMN was detected by autoradiography (bottom).

To evaluate what fraction of hnRNP A1 bound to U7 snRNA associates with the ternary complex, GST-PTBP1 and U7 snRNA alone or their complex were incubated with a HeLa nuclear extract, the samples split in two halves and incubated with either SA (Fig. 7B, lane 1-3) or GSH beads (Fig. 7B, lane 4-6), via U7 snRNA and recombinant PTBP1, respectively. Bound proteins were analyzed by silver staining (top) and Western blotting (bottom). Compared to the amount of hnRNP A1 bound to U7 snRNA and purified on SA beads, only a small fraction was detected in the ternary complex immobilized on GSH beads (Fig. 7B, top and bottom, compare lanes 3 and 6). Interestingly, the association of Matrin3 with recombinant PTBP1 was clearly stimulated in the presence of U7 snRNA (Fig. 7B, top and bottom, compare lanes 4 and 6) (see below).

We also tested whether hnRNP A1 forms a ternary complex with PTBP2 bound to U7 snRNA. In addition, to determine which protein bands in the silver-stained gel are contributed solely by recombinant proteins, GST tagged PTBP1 and PTBP2 were immobilized on GSH beads in the presence of a buffer rather than nuclear extract (Fig. 7C, lanes 4 and 7). Each of the two GST-tagged proteins associated with hnRNP A1 only when pre-bound to U7 snRNA (Fig. 7C, compare lanes 2 and 3, and lanes 5 and 6). The same results were obtained with cytoplasmic extracts, although since these extracts contain lower levels of endogenous hnRNP A1, the formation of a ternary complex was less efficient. Altogether, these results suggest that PTBP1 (and its paralogues) and hnRNP A1 (and its paralogues) are capable of forming a ternary complex with U7 snRNA that in turn associates with SMN, a subunit of the SMN complex.

In the ternary complex, the amount of SMN strictly correlated with the amount of hnRNP A1. To test whether hnRNP A1 directly recruits SMN, GST alone or GST fused to hnRNP A1 were incubated with *in vitro* synthesized ^35^S-labelled SMN and immobilized on glutathione beads. GST protein failed to detectably interact labeled SMN (Fig. 7D, lane 2). Importantly, as much as ∼20% of the labeled proteins used in the binding assay was recovered via GST-tagged hnRNP A1 (Fig. 7D, lane 3). We conclude that hnRNP A1 directly interacts with SMN and recruit this subunit of the SMN complex to U7 snRNA or a ternary complex additionally containing PTBP1.

### U7 snRNA stabilizes complexes of PTBP1 with its binding partners containing PRI motifs

One interesting and largely unexplored aspect of the PTBP biology is that it forms complexes with proteins containing a single or multiple copies of PTBP RRM2 Interacting (PRI) motif with the (S/G)(I/L)LGxxP consensus. This motif interacts with the helical face of the RRM2 that does not participate in RNA recognition, allowing formation of ternary complexes in which PTBP1 is bound to both its RNA target and a PRI-containing protein. These proteins include Matrin3, Raver1, Raver2, CCAR1 and CCAR2 (57–61).

We tested several cytoplasmic and nuclear extracts from HeLa cells for the presence of PRI-containing proteins and to determine how U7 snRNA affects binding of these proteins to PTBP1. In these experiments, U7/WT RNA (or is mutants) and PTBP individually or together were incubated with the extracts and bound complexes captured on streptavidin (SA) beads via biotin attached to the RNA, or on glutathione (GSH) beads via GST tag attached to the recombinant PTBP1, as described for Fig. 7B. In the initial experiment, we used U7/WT either alone or in a complex with GST-PTBP1, and RNA-containing complexes formed in a nuclear extract were immobilized on SA beads. To determine which protein bands in silver-stained gels are contributed by the recombinant protein, a complex of GST-PTBP1 and U7/WT was bound to SA beads in the presence of a buffer (Fig. 8A, top, right lane). In agreement with the results shown above, the material bound to U7/WT contained both isoforms of PTBP and its interacting partner with a PRI motif, Matrin3. Note that Matrin3 co-migrates with other proteins, yielding a predominant silver-stained band migrating just below the 130 kDa size marker (Fig. 8A, top, lane 2). Pre-incubation of U7/WT RNA with GST-PTBP1 to form their complex did not change the amount of endogenous PTBP1 immobilized on SA beads, indicative of excess of RNA used in the experiment (Fig. 8A, top, lane 3). Surprisingly, the material bound by this complex contained a protein of 130 kDa, which was missing in lane containing material bound to U7/WT RNA alone. This band was identified by mass spectrometry and Western blotting as a different PRI motif-containing protein, CCAR2 (Fig. 8A, top and bottom, compare lanes 2 and 3, arrow). Only a small amount of this protein was detected by Western blotting in the sample lacking recombinant GST-PTBP1, suggesting that CCAR2 preferentially interacts with recombinant PTBP1, hence behaves in a sharp contrast to Matrin3, which co-purifies with endogenous PTBP1 (Fig. 7B).

**Fig. 8.**
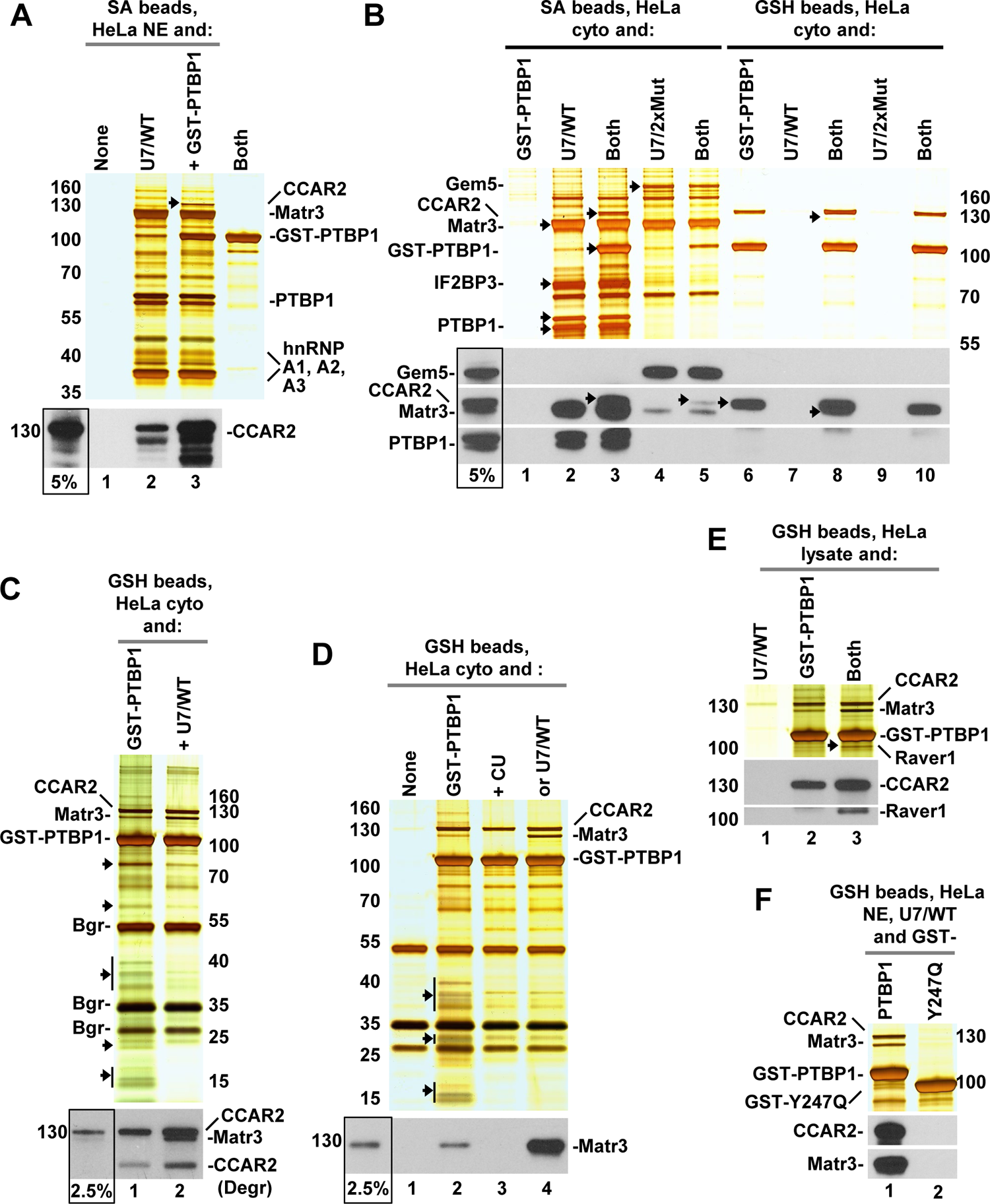
U7 snRNA stabilizes complexes of PTBP1 with PRI motif-containing proteins. **A.** Affinity purification of proteins from a HeLa nuclear extract (NE) bound to U7/WT RNA alone (lane 2) or in complex with recombinant GST-PTBP1 (lane 3). The bound proteins were immobilized on SA beads and detected by silver staining (top). CCAR2 band in lane 2 is indicated with an arrow (top). The presence of CCAR2 was monitored by Western blotting (bottom). In the bottom panel, the input lane (boxed) contains ∼5% of the nuclear extract used in the binding experiment. In the last lane of silver-stained gel (top), the complex of U7/WT RNA and GST-PTBP1 was immobilized on SA beads in the presence of a buffer instead of nuclear extract. **B.** Affinity purification of proteins from a HeLa cytoplasmic extract bound to U7/WT or U7/2xMut RNAs alone or in complex with recombinant GST-PTBP1. Bound proteins were immobilized on SA (lanes 1-5) or GSH beads (lanes 6-10) and detected by silver staining (top) or Western blotting (bottom). In the silver-stained gel, the arrows indicate bands containing Gemin5, CCAR2, Matrin3, PTBP1, IGF2BP3 and recombinant GST-PTBP1. In the middle Western blotting, CCAR2 and Matrin3 were detected simultaneously by specific antibodies, and the arrows indicate the top band corresponding to CCAR2 (lanes 3, 5 and 6), and bottom band corresponding to Matrin3 (lane 8). **C.** Recombinant GST-tagged PTBP1 alone (lane 1) or pre-bound to U7 snRNA (lane 2) were incubated with a HeLa cytoplasmic extract and proteins bound to GST-PTBP1 were immobilized on glutathione (GSH) beads and analyzed by silver staining (top) or Western blotting (bottom). In silver-stained gel, the arrows indicate multiple ribosomal proteins that were eliminated in the presence of U7/WT RNA (lane 2). Background bands (Bgr) represent proteins non-specifically bound to GSH beads (see also lane 1 in Fig. 7A and C). In the bottom panel, the input lane contains ∼2.5% of the material used in the binding experiment. A degradation byproduct of CCAR2 (Degr) is indicated. **D.** Recombinant GST-tagged PTBP1 alone (lane 2) or in complex with CU oligonucleotide (lane 3) or WT/U7 RNA (lane 4) were immobilized on GSH beads, washed extensively, and incubated with a HeLa cytoplasmic extract. The bound material was analyzed by silver staining (top) or Western blotting (bottom). Lane 1 contains proteins that non-specifically bind to GSH beads in the absence of recombinant GST-PTBP1. The arrows indicate ribosomal proteins (lane 2). **E.** Recombinant GST-tagged PTBP1 alone (lane 2) or pre-bound to U7/WT RNA (lane 3) were incubated with a lysate from HeLa cells and proteins bound to GST-PTBP1 were immobilized on glutathione (GSH) beads and analyzed by silver staining (top) or Western blotting (bottom). **F.** Recombinant GST-tagged PTBP1 or its Y247Q mutant, as indicated, were both pre-bound to U7/WT RNA, incubated with a HeLa nuclear extract (NE) and immobilized on GSH beads. The bound material was analyzed by silver staining (top) or Western blotting (bottom).

To confirm this difference in the behavior of Matrin3 and CCAR2, we extended the experiment by using a HeLa cytoplasmic extract and two RNAs, U7/WT and U7/2xMut, that drastically differ in their ability to bind PTBP1. As determined by capturing RNA-containing complexes on SA beads, U7/WT, but not U7/2xMut RNA, readily interacted with PTBP1 (Fig. 8B, top and bottom, compare lanes 2 and 4). The same binding pattern was observed for Matrin3 (Fig. 8B, bottom, compare lanes 2 and 4). Note that the difference in the amount of Matrin3 bound to U7/WT and U7/2xMut RNAs can be visualized by Western blotting, but not by silver staining, as Matrin3 co-migrates with hnRNP U and U-like 1 that interact with both RNAs. As expected, U7/2xMut bound endogenous Gemin5 (Fig. 8B, top and bottom, lane 4). The material bound by a complex of U7/WT RNA and GST-PTBP1 contained readily detectable amounts of CCAR2 (Fig. 8B, top and bottom, compare lanes 2 and 3, arrow). Only a residual amount of this protein was detected by Western blotting in the material purified by the complex of U7/2xMut and GST-PTBP1, in a perfect correlation with the small amount of GST-PTBP1 (Fig. 8B, top and bottom, compare lane 5, arrow).

Second half of each sample was in parallel incubated with GSH beads, allowing for the selection of proteins that bind recombinant PTBP1 either alone or in complex with U7 snRNA. In all samples containing GST-tagged PTBP1, regardless of whether the RNA was added or not, CCAR2 was readily detectable by silver staining and its presence confirmed by Western blotting (Fig. 8B, top and bottom, lanes 6, 8 and 10). This result demonstrates that CCAR2 binds recombinant PTBP1 in an RNA-independent manner, although U7/WT RNA, but not U7/2xMut RNA, has a clear stimulatory effect on this interaction, resulting in more CCAR2 purified on GSH beads (Fig. 8B, top and bottom, compare lane 8 with lanes 6 and 10). Both silver staining and Western blotting failed to detect Matrin3 in the material immobilized on GSH beads when just GST-PTBP1 was used, although a small amount of this protein was detected in the presence of U7/WT RNA (Fig. 8B, top and bottom, compare lanes 6 and 8, arrows). U7/2xMut RNA, which does not bind PTBP1, had no effect (Fig. 8B, lane 10). We conclude that in contrast to Matrin3, which preferentially interacts with endogenous PTBP1, CCAR2 exists in HeLa cytoplasmic extracts predominantly as a free protein, available to from a complex with recombinant PTBP1.

To confirm that U7 snRNA stimulates binding of recombinant PTBP1 to Matrin3 and CCAR2, we repeated this experiment with a HeLa cytoplasmic extract and increased the intensity of silver staining to detect less abundant proteins immobilized on GSH beads. Again, GST-PTBP1 in the absence of U7/WT RNA interacted with only a small amount of Matrin3 (Fig. 8C, lane 1), consistent with this protein being primarily associated with endogenous PTBP1. In the presence of U7/WT RNA, this amount clearly increased, as determined by both silver staining and Western blotting (Fig. 8C, top and bottom, lane 2), suggesting that U7 snRNA either stimulates formation of the complex between recombinant PTBP1 and Matrin3, or stabilizes this complex. The same stimulatory effect of U7 snRNA was also observed on binding CCAR2. Interestingly, the U7 snRNA-mediated increase in the amount of Matrin3 and CCAR2 bound to recombinant PTBP1 was accompanied by a disappearance of multiple bands that were purified in the absence of U7 snRNA (Fig. 8C, lane 1, arrows). Mass spectrometry identified these bands as ribosomal proteins of the large and small subunits.

We tested whether the opposing effects of U7 snRNA on the ability of recombinant PTBP1 to interact with ribosomal proteins and the two PRI proteins, Matrin3 and CCAR2, can be reproduced using any RNA that is recognized by PTBP1. In this experiment, GST-PTBP1 was prebound to either U7/WT RNA or CU oligonucleotide, and the resultant complexes were immobilized on GSH beads and incubated with a Hela cytoplasmic extract. Strikingly, pre-binding of CU oligonucleotide to GST-PTBP1 reduced the amount of not only ribosomal proteins but also Matrin3 and CCAR2 (Fig. 8D, lane 3). In contrast, pre-binding of U7/WT RNA to GST-PTBP1, while eliminating ribosomal proteins, dramatically stimulated the interaction of GST-PTBP1 with Matrin3 (Fig. 8D, lane 4). The stimulatory effect of U7 snRNA on the interaction between CCAR2 and recombinant PTBP1 was less evident in this experiment, largely due to relatively strong silvers staining of CCAR2 in lane 2.

Only a limited number of HeLa nuclear extracts and whole cell lysate allowed for a successful co-purification of Raver1, another PTBP1 binding partner containing a PRI motif. Binding of also this protein, largely or exclusively of nuclear localization, to recombinant PTBP1 was clearly stimulated by U7 snRNA, thus mimicking the behavior of Matrin3 and CCRA2 (Fig. 8E, top and bottom, compare lanes 2 and 3).

Matrin3, CCAR2, Raver1 and other proteins containing PRI motifs interact with the helical face of RRM2, with the key role being played by tyrosine 247 and its substitution with Q is sufficient to disrupt the interaction (59). We expressed GST-tagged PTBP1 containing this mutation (see Fig. 3C) and tested its effect on the interaction with PRI motif-containing proteins in the presence of U7/WT RNA. As expected, while GST-PTBP1 interacted with easily detectable amounts of Matrin3 and CCAR2 (Fig. 8F, top and bottom, lane 1), the Y247Q mutant was fully impaired in binding these two proteins (Fig. 8F, top and bottom, lane 2).

## DISCUSSION

The U1, U2, U4, U4_atac_, U5, U11 and U12 spliceosomal snRNAs are generated by RNA polymerase II and they all contain an Sm site, a conserved 9-nucleotide sequence with the AAUUU(U/G)UGG consensus (18,91,92). *In vivo*, snRNAs containing this sequence are recognized by Gemin5 (24,26), which subsequently delivers them to the remaining subunits of the SMN complex for a multi-step and still poorly understood process of assembling a heptameric ring around the Sm site. The spliceosomal Sm ring consists of Sm subunits D1, D2, B, D3, E, F and G (10,18,93). Three of them, SmD1, SmB and SmD3, contain within their C-terminal regions multiple symmetrically dimethylated arginines, which are recognized by the SMN protein, hence facilitating the stepwise assembly process (15). The resultant RNA/protein complexes, referred to as snRNP cores, each consisting of one spliceosomal snRNA and an identical heptameric ring undergo multiple maturation steps in the cytoplasm and the nucleus, including the recruitment of snRNP-specific proteins, ultimately becoming fully functional spliceosomal snRNPs and the key constituents of major and minor spliceosomes (93).

Animal cells additionally contain U7 snRNP, a distant relative of the spliceosomal snRNPs that has no role in splicing and instead functions as an RNA-guided endonuclease and 5’-3 exonuclease in 3’ end processing of replication-dependent histone pre-mRNAs (3,4). The Sm site in U7 snRNA with the consensus of AAUUUGUCUAG departs from the spliceosomal consensus in both the sequence and length and nucleates the assembly of a different ring in which SmD1 and D2 are replaced by related Lsm10 and Lms11 (1,9,15). The unusual Sm site in U7 snRNA is not recognized by Gemin5 (24), although at least some other components of the SMN complex, including the SMN subunit, are required for more downstream steps in the assembly of the U7-specific Sm ring (9,45,46). The identity of the protein(s) that recognizes distinct features of the Sm site in U7 snRNA and substitutes for Gemin5, hence promoting the incorporation of Lsm10 and Lsm11 into the Sm ring of U7 snRNP has not been previously determined.

### Binding of PTBP1 to U7 snRNA

Using RNA affinity chromatography and extracts from mammalian cells, we identified PTBP1 as a protein that recognizes the U7-specific Sm site. A 69-nucleotide U7 snRNA with the wild-type sequence of its Sm site forms a tight complex not only with endogenous PTBP1 but also with bacterially expressed recombinant PTBP1 and PTBP1 generated by *in vitro* translation. The binding of PTBP1 to U7 snRNA is severely inhibited by changing its Sm site to the spliceosomal consensus. In cytoplasmic extracts, this mutant RNA, while failing to interact with PTBP1, forms a complex with Gemin5, in agreement with previous *in vivo* studies where the same U7 snRNA mutant promoted the assembly of an Sm ring with SmD1 and SmD2 (76,77). Binding of PTBP1 to U7 snRNA also requires an upstream sequence, CUCUU, that base-pairs with histone pre-mRNA, defining the substrate specificity of U7 snRNP. Thus, PTBP1 simultaneously recognizes two functionally essential sequences in U7 snRNA, the substrate base-pairing (SBP) site and the Sm binding site, arguing that it may act *in vivo* as the functional counterpart of Gemin5.

The interaction of PTBP1 with the SBP and Sm sites is consistent with the known preference of PTBP1 for CU tracts (50,94). The CU dinucleotides are the hallmark of both sites in U snRNA and they are invariably conserved in all known and presumptive metazoan U7 snRNA sequences (95), being at the same time absent from the Sm site in spliceosomal snRNAs (10,96). Human U7 snRNA contains two additional CU-containing elements, one located in the loop (nucleotides 44-49), and the other one located near the 3’ end (nucleotides 62-66). At least the 3’ terminal CUCUU sequence contributes to the interaction between PTBP1 and U7 snRNA. Mature U7 snRNA lacks part of this sequence as a result of 3’ end trimming that occurs after the completion of the Sm ring. This shorter U7 snRNA forms a significantly less stable complex with PTBP1 in a mobility shift assay (not shown).

The importance of the CUCUUU substrate base pairing site in U7 snRNA for binding PTBP1 may explain the usual evolutionary conservation of this sequence, which is virtually invariable throughout all metazoans (95). The CUCUUU sequence base pairs with the AAAGAG purine core of the HDE in histone pre-mRNAs. In many histone pre-mRNAs, the HDE sequence departs from this consensus, resulting in one or more mismatches in the duplex formed with the U7 snRNA (8). It remains puzzling what mechanisms have prevented co-evolution of the two CUCUUU sequence in U7 snRNA and the AAAGAG HDE in histone pre-mRNAs into mixed purine/pyrimidine tracts. Studies with *in vitro* assembled semi-recombinant (97) and fully recombinant U7 snRNP (3,98) demonstrated that extensive compensatory mutations within the base pairing regions, including switching all six pyrimidines with purines in U7 snRNA do not affect the function of U7 snRNP *in vitro*, which efficiently cleaves histone pre-mRNA containing all six purines substituted with pyrimidines. Interestingly, the same mutant U7 snRNA when expressed in vivo yielded nuclear extracts poorly active in processing (8), a potential indication of the importance of the CUCUUU sequence. Clearly, the recognition of the CUCUUU site by PTBP1 as an essential step in the biogenesis of U7 snRNP would place a strong evolutionary constraint on this sequence, preventing its major sequence variations. PTBP1 is a versatile and highly abundant RNA binding protein, regulating a plethora of processes, including alternative splicing, cap-independent translation, 3’ end processing, mRNA degradation and mRNA localization (63,99). In human cells, PTBP1 has two paralogs, PTBP2 and PTBP3. These three highly related proteins are believed to fulfill both specific and redundant functions, and our studies demonstrated that PTBP1 and PTBP2 are equally efficient in binding U7 snRNA, and PTBP3 may bind with lower affinity (data not shown). PTBP2 is neuron-specific and plays a key role in converting constantly cycling neuronal progenitor cells into terminally differentiated and mitotically inactive neuronal cells (67,100). The function of PTBP3 is largely unknow but it may be specifically linked to the development of hematopoietic cell linkages (67,99).

PTBP1 and its paralogues contain four RRMs, and the two N-terminal domains function independently, binding relatively short CU stretches often located in loops (50,52). In contrast, RRM3 and RRM4 tightly interact through multiple contacts, likely forming a single functional unit in which the two RRMs have a fixed orientation relative to each other (50,51,101). They can simultaneously bind two CU-containing motifs in antiparallel orientation if they are separated by ∼15 nucleotides, resulting in RNA looping (50,102). The spacer between the CUCU and UCU motifs in the SBP and Sm sites in U7 snRNA has 11 nucleotides and is likely sufficiently long to be looped out following the recognition of these site by the RRM34 tandem. RRM1 and RRM2 may in turn interact with the two remaining CU-containing motifs, one in the 3’ terminal loop, and the other near the 3’ end. This may result in a very tight RNA/protein complex, with both components adopting a unique conformation and spatial arrangement, explaining inaccessibility of PTBP1 in the complex for anti-PTBP1 antibodies. Clearly, structural studies are required to determine how PTBP1 and its paralogues interact with U7 snRNA.

### Binding of IGF2BP3 to U7 snRNA

In addition to PTBP1, our biochemical assays with Hela cytoplasmic extracts identified IGF2BP3 as a protein that interacts with U7 snRNA in a manner that requires its unique Sm site. Strikingly, IGF2BP3 also requires the CUCU sequence in the site that base-pairs with histone pre-mRNA, thus mimicking the binding specificity of PTBP1. Surprisingly, binding of IGF2BP3 to U7 snRNA was not inhibited by excess of the CU oligonucleotide, strongly contrasting with PTBP1. It is possible that IGF2BP3 recognizes certain structural features within U7 snRNA that depend on the wild type sequences of the SBP and Sm sites.

Binding of IGF2BP3 to U7 snRNA was detected in cytoplasmic extracts from HeLa cells but not in lysates from mouse C2C12 skeletal muscle cells and human SH-SY5Y neuroblastoma cells and could not be reproduced using *in vitro* generated IGF2BP3. These results suggest that IGF2BP3 may require another protein or a posttranslational modification for binding to U7 snRNA. Intriguingly, binding of IGF2BP3 to U7 snRNA is affected by the presence of EDTA, suggesting that magnesium or other metal ions play an important role in IGF2BP3 directly binding to the RNA, to RNA-bound protein(s) or/and appropriate RNA folding. In addition, IGF2BP3 has an increased affinity to U7 snRNA containing phosphate at the 5’ end, indicative of a preference for the presence of a 5’ cap (103). The importance of these observations for the biogenesis of U7 snRNP, if any, and potentially for other cellular functions of U7 snRNA remains unknown.

### Binding of hnRNP A1 to U7 snRNA

In addition to PTBP1 and IGF2BP3, affinity purification identified hnRNP A1 as a third protein that interacts with human U7 snRNA. hnRNP A1 has a strong specificity for UAG sequences (55,56,90), and two of them are found in U7 snRNA, with one separating the HDE- and Sm-binding sites and the other being an integral part of the Sm-binding site (95). Both these sequences are universally conserved in vertebrate U7 snRNAs, but absent from the *Drosophila* U7 snRNA, which in these positions contains GA and UG dinucleotides. hnRNP A1 contains two RRMs and they also interact with each other forming a single binding platform capable of interacting with bipartite binding sites in RNAs (55,104). These two RRMs likely act in conjunction to recognize the two UAG sites in U7 snRNA, although structural studies are required to substantiate this prediction.

In contrast to PTBP1 and IGF2BP3, hnRNP A1 readily interacted with the U7 snRNA SPL mutant in which the Sm site was mutated from AAUUUGUCUAG to the spliceosomal consensus AAUUU<u>U</u>U<u>GG</u>AG (changed nucleotides are underlined), initially disqualifying this proteins as a U7-specific counterpart of Gemin5. While hnRNP A1 preferentially binds UAG motifs, it makes key contacts with the AG dinucleotide (50,55). This dinucleotide was left unchanged in the SPL mutant, creating together with the upstream UAG trinucleotide a sufficiently strong binding site for hnRNP A1. Substituting the two AG dinucleotides with GA, a mutation that maintain the overall nucleotide composition of U7 snRNA, abolished hnRNP A1 binding.

### PTBP1 and hnRNP A1 form a ternary complex with U7 snRNA: a role in the assembly of U7-specific ring?

Our biochemical studies demonstrated that PTBP1 and hnRNP A1 can interact with U7 snRNA both individually, forming two different binary complexes, and simultaneously, forming a ternary complex. This is reminiscent of SLBP and 3’hExo, the two proteins that bind either alone or together to the terminal stem-loop structure of replication-dependent histone mRNAs (105–107). Binding of PTBP1 and hnRNP A1 to the relatively short region of U7 snRNA of only 20 nucleotides encompassing the SBP and Sm sites is surprising and may be indicative of an important biological function. The close association of SLBP and 3’hExo at the end of replication-dependent histone mRNAs was later shown to be essential for selective degradation of these mRNAs upon the completion or interruption of DNA replication (108,109).

PTBP1 and hnRNP A1 do not appear to cooperate in recognizing their targets in U7 snRNA and forming a ternary complex. On the contrary, the elimination of binding sites for one protein, improves binding of the other protein, suggesting that they compete rather than cooperate in binding U7 snRNA, and this competition likely results from the proximity of the PTBP1- and hnRNP A1-bindning sites. Indeed, a relatively small fraction of hnRNP A1 found in a binary complex with U7 snRNA forms a ternary complex additionally containing PTBP1. It is possible that the formation of a ternary complex containing PTBP1 and hnRNP A1 requires a particulate conformation of U7 snRNA that exposes target sites for each of the two proteins and accommodates both proteins on a relatively short RNA fragment. *In vivo* this effect may be achieved by a sequential order of events, with PTBP binding first, followed by hnRNP A1, or vice versa. Interestingly, the upstream UAG binding target for hnRNP A1 is located within the spacer between the SBP and Sm sites that might be looped out following binding of PTBP1 to become available for hnRNP A1 binding. The second UAG motif is located at the 3’ end of the Sm site, immediately downstream of the presumptive PTBP1-binding motif, UCU. We speculate that this direct juxtaposition of PTBP1 and hnRNP A1 binding sites, which are at the same time the key specificity determinants of the Sm site in U7 snRNA, may be important for subsequent binding of the seven subunits of the Sm ring, including Lsm10 and Lsm11.

Two observations support a role for PTBP1 and/or hnRNP A1 in the assembly of the Sm ring on U7 snRNA. First, the 3’ region of the U7-specific Sm site, <u>UC</u>U<u>AG,</u> is absolutely conserved in all known U7 snRNA sequences in vertebrates (95). Of this short motif, four nucleotides (underlined) are essential for the formation of U7 snRNP *in vivo*, and their individual substitutions with any other nucleotide prevents the assembly of both U7 snRNP and spliceosomal snRNPs (110). These results indicate that individual substitutions of the four essential nucleotides, while abolishing important interactions required for the biogenesis of U7 snRNP, generate at the same time sequences that have no affinity for Gemin5. The effect of these mutations is therefore similar to the effect of the U7/2xGA mutation tested in our studies and suggests that binding of Gemin5 may require additional substitutions that make the U7-specific Sm site more similar to the spliceosomal Sm site.

The second observation is that hnRNP A1 directly interacts with SMN, delivering this key subunit of the SMN complex to both the hnRNP A1/U7 snRNA binary complex and to a ternary complex additionally containing PTBP1. SMN was previously shown to interact with a plethora of RNA binding proteins, including hnRNP R and hnRNP Q (42,111). We anticipate that in addition to SMN, at least some other subunits of the SMN complex are required for the assembly of U7 snRNP, and they may be recruited to U7 snRNA during later steps in the assembly process. Further studies are required to investigate this and other possibilities.

### Binding of Gemin5 to U7 snRNA

The recognition of the spliceosomal AAUUUUGG consensus sequence by Gemin5 (24–26) critically depends on the second adenosine, and the first and third uridines (24,86–89). Surprisingly, despite containing the same nucleotides in these positions, U7 snRNA does not interact with Gemin5 both *in vivo* (24) and *in vitro* (this study). Metazoans likely developed efficient safeguard mechanisms to prevent binding of Gemin5 to the U7-specific Sm site that could potentially result in the assembly of an erroneous ring on U7 snRNA with SmD1 and SmD2 instead of Lsm10 and Lsm11. Clearly, the fact that SmD1 and SmD2 are ∼500-fold more abundant than Lsm10 and Lsm11 increases the risk of generating an entirely spliceosomal ring around U7 snRNA and a functionally defective and potentially harmful U7 snRNP variant.

Our studies provide potential clues to understanding how some of these safeguard mechanisms might work. In both RNA-mediated affinity purification and EMSA experiments, U7 snRNA with the wild type Sm site failed to detectably interact with Gemin5, but a visible increase was observed in the presence of short oligonucleotides capable of sequestering PTBP1 or hnRNP A1. Thus, Gemin5 is unable to compete with PTBP1 and hnRNP A1, which have higher affinities for the U7-specific Sm site and are more abundant. The nuclear fraction of hnRNP A1 is more efficient than the cytoplasmic fraction in forming a ternary complex with U7 snRNA and PTBP1. Plausibly, the two proteins may bind U7 snRNA co-transcriptionally in the nucleus, where Gemin5 is limiting, and assist U7 snRNA to the cytoplasm, effectively shielding the U7-specific Sm site against much higher Gemin5 concentrations in this compartment.

An improved binding of Gemin5 to the U7-specific Sm site was also observed in the presence of an oligonucleotide complementary to the first 18 nucleotides of the U7 snRNA. By forming a duplex structure, this oligonucleotide eliminated the upstream site for PTBP1, but also likely affected overall folding of U7 snRNA, resulting in a substantial increase in the amount of bound Gemin5. Structural studies on recombinant U7 snRNP revealed that the C and G nucleotides within the UCUAG sequence at the 3’ end of the Sm site in U7 snRNA form an intramolecular base pair (3). These two nucleotides are specific to the Sm site in U7 snRNA and according to our current studies are recognized by PTBP1 and hnRNP A1, respectively. Further studies are required to determine whether the CG base pair is an intrinsic feature of U7 snRNA structure in a free state. If so, it may affect the repertoire of proteins that bind U7 snRNA and/or modify the mode of their interactions. Clearly, other possible structural determinants within the Sm site, including long range contacts, may limit binding of Gemin5 and instead favor interactions with RNA binding proteins that participate in the U7-specific Sm ring assembly pathway.

### U7 snRNA stabilizes the interaction between PTBP1 and its binding partners

PTBP1 uses its RRM2 to interact with a group of proteins containing one or more copies of the PTBP RRM2 Interacting (PRI) motif with the (S/G)(I/L)LGxxP consensus (57–61). These proteins include Matrin3, Raver1, its paralogue Raver2, CCAR1 and its paralogue CCAR2. The PRI motif interacts with the helical face of the RRM2 that does not participate in RNA recognition, allowing formation of ternary complexes with PTBP1 bound to both its RNA targets and PRI proteins.

Our binding experiments with full length PTBP1 and various batches of HeLa cytoplasmic, nuclear and whole cell extracts resulted in co-purification of all five PTBP interacting proteins. However, their presence depended on the batch of HeLa cells used to prepare the extracts. While Matrin3 universally copurified with PTBP1 and was present in both the cytoplasmic and nuclear extracts, Raver1-2 and CCAR1-2 were detected only occasionally, with Raver1-2 displaying apparently nuclear distribution. Interestingly, as demonstrated by U7 snRNA-mediated purification, Matrin3 was almost exclusively associated with endogenous PTBP1, contrasting with both Raver1-2 and CCAR1-2, which likely existed in a free state available for binding to recombinant PTBP. These observations suggest that the formation of complexes between PTBP1 and its binding partners may be dynamically regulated in cells, depending on their type and physiological status. Intriguingly, the interaction of PTBP1 with its binding partners is stabilized in the presence of U7 snRNA. In contrast, a different PTBP1 RNA target containing multiple CU dinucleotides resulted in destabilization of the same complexes. The importance of this findings is currently unclear but is supports a functional relationship between PTBP1 and U7 snRNA.

Matrin3, Raver1, and likely other proteins containing PRI motifs assist PTBP1 in regulating alternative splicing of multiple transcripts (58,112–114). The close partnership between PTBP1 and the PRI proteins may also serve other purposes, including the recognition of internal ribosome entry segments (IRES) in viral and cellular mRNAs (115) and 3’ end processing (57). Interestingly, in our affinity purification experiments, recombinant GST-tagged PTBP1 interacted with ribosomal proteins from cytoplasmic extracts, potentially reflecting the role of PTBP1 in cap-independent translation.

Raver1 was initially identified as a dual compartment protein, residing primarily in the nucleus, but being redistributed to sites of cytoskeletal assembly in differentiating muscle cells (116–118). A similar redistribution from the nucleus was also demonstrated for PTBP1, which migrates to cytoplasmic protrusions and focal adhesions, targeting to these sites various transcripts for local translation that may control cell migration and attachment (119). It is possible that in this translocation between various cellular compartments PTBP1 and Raver1 closely co-operate with each other forming larger complexes with additional proteins, including cytoskeletal components, and a plethora of mRNAs that bind to multiple RNA binding domains in the two proteins (99). It is tempting to speculate that the stabilizing effect of U7 snRNA on the interaction between PTBP1 and Raver1 and other PRI proteins may in at least some cellular settings facilitate formation of these larger complexes and regulate their destination and function.

## Conclusions

Our biochemical studies identified PTBP1 and hnRNP A1 as proteins that recognize unique features of the U7-specific Sm site, making additional contacts with other regions of U7 snRNAs. Further studies are required to determine whether these two proteins, either separately or together, function in the assembly of the U7-specific Sm ring containing Lsm10 and Lsm11. The only result directly supporting this possibility is that U7 snRNA forms a larger complex containing PTBP1, hnRNP A1 and SMN, likely directly recruited to the U7 snRNA by hnRNP A1.

In human cells PTBP1 has two paralogues, PTBP2 and PTBP3, that share as much as 75% sequence identity that likely play both redundant and specific roles. PTBP2 is mostly expressed in neuronal cells and is one of the key players in differentiating neuronal progenitor cells into mature neurons (67,100). U7 snRNA interacts with the two paralogues that are expressed in a tissue-specific manner during cell differentiation and stabilizes complexes of PTBP with its binding partners Matrin3, Raver1/2 and CCAR1/2. One attractive possibility is that U7 snRNA, and potentially other U7 snRNP components, become repurposed for new roles in post-mitotic cells that cease to express replication-dependent histone pre-mRNAs.

## MATERIALS and METHODS

### Antibodies

The following commercial antibodies were used in this study: anti-Matrin3 (A300-591A, BETHYL), anti-CCAR2 (22638-1-AP, Proteintech), anti-IGF2BP3 (PA5-46704, Invitrogen), anti-PTB1 (12582-1-AP, Invitrogen), anti-PTB2 (55186-1-AP, Proteintech), anti-Raver1 (A303-939A, BETHYL), anti-hnRNP A1 (11176-1-AP, Proteintech), anti-hnRNP U1 (MA5-35471, Invitrogen), anti-hnRNP U like 1, (10578-1-AP, Proteintech), anti-hnRNP A2 (sc-374053, Santa Cruz), anti-SMN (60154-1-Ig, Proteintech), anti-Gemin2 (2E17, SIGMA), anti-Gemin3 (12H12, sc-57007, Santa Cruz), anti-Gemin4 (17D10, sc-136199, Santa Cruz), anti-Gemin5 (A301-325A, BETHYL, anti-Gemin5 (10G11, Santa Cruz).

### RNAs

The following RNAs synthesized by Dharmacon (Lafayette, CO, USA) were used in this study. All sequences are written in 5′-3′ orientation.

U7/63mer: CAGTGTTACAGCTCTTTTAGAATTTGTCTAGTAGGCTTTCTGGCTTTTTACCGGAAAGCCCCT

U7/WT: CAGUGUUACAG<u>CUCUUU</u>UAG<u>AAUUUGUCUAG</u>UAGGCUUUCUGGCUUUUUA

CCGGAAAGCCCCUCUUAUG U7/SPL:

CAGUGUUACAGCUCUUUUAGAAUUU**<u>U</u>**U**<u>GG</u>**AGUAGGCUUUCUGGCUUUUUA

CCGGAAAGCCCCUCUUAUG-18S-3’Biot U7/PU:

CAGUGUUACAG**<u>GAGA</u>**UUUAGAAUUUGUCUAGUAGGCUUUCUGGCUUUUUA

CCGGAAAGCCCCUCUUAUG-18S-3’Biot

U7/2xMut: CAGUGUUACAG**GAGA**UUUAGAAUUU**<u>U</u>**U**<u>GG</u>**AGUAGGCUUUCUGGCUUUUU

ACCGGAAAGCCCCUCUUAUG-18S-3’Biot U7/2xGA:

CAGUGUUACAGCUCUUUU**<u>GA</u>**AAUUUGUCU**<u>GA</u>**UAGGCUUUCUGGCUUUUUA

CCGGAAAGCCCCUCUUAUG-18S-3’Biot

3’BiotDmU7: UGAAAAUUUUUAU<u>UCUCUUU</u>GA<u>AAUUUGUCUUG</u>GUGGGACCCUUUGUCUA

GGCAUUGAGUGUUCCCGUU-18S-3’Biot Anti-U7: mAmAmAmGCUmAmGmCmUmGmUmAmAmCmAmCmUmU (18-mer) U7 5’ end: CAGUGUUACAGCUCUUUUAG (20-mer)

CU: 5’ CUCUCUCUCUUUCUUU 3’ (16-mer) UAG: UAUGAUAGGGACUUAGGGUG (20-mer)

### Cell culture and preparation of cytoplasmic and nuclear extracts

HeLa cells were grown by Cell Culture Company (Minneapolis, MN), shipped on dry ice as a frozen pellet and used after thawing. Nuclear extracts were prepared, as described (74,120,121). The crude cytoplasm generated after collecting HeLa nuclei was supplemented depending on experiments with NP40 to a final concentration ranging from 0.025% to 0.1% and used to prepare cytoplasmic extracts, as described (122), with the ultracentrifugation and dialysis steps being omitted.

### Preparation of whole cell lysates

Pellets of HeLa, C2C12 and SH-SY5Y cells were resuspended in 0.5% NP40 lysis buffer (40 mM Tris pH 7.5, 150 mM NaCl, 0.5% NP40), using 5 ml of buffer per 1 ml of cell pellet. The cell suspensions were rotated in cold room for 30 min and spun for 30 min at 30,000 x g in a refrigerated centrifuge.

### Isolation and identification of mammalian proteins interacting with U7 snRNA

Proteins that bind U7/WT RNA and its mutants were purified essentially, as previously described (74,75,123). Briefly, a HeLa cytoplasmic extract (750 µl) was mixed with U7/WT snRNA or its mutants (0.75-7.5 µg, depending on experiment) and rotated in cold room for 60 min. The sample was spun in cold room for 10 min at 13,500 x g and the supernatant loaded over 25 µl of streptavidin beads (Sigma) and rotated in cold room for 1 hr. The beads were gently spun, rinsed twice and rotated for 60 min with a buffer containing 40 mM HEPES pH 7.9, 150 mM NaCl, 4.5 mM MgCl_2_, 0.5 mM DTT and 0,025% NP40.

A fraction of each sample was separated by electrophoresis in a 4-12% SDS/polyacrylamide gel (SDS/PAGE) and the bound proteins visualized by silver staining and identified by mass spectrometry in the Laboratory of Mass Spectrometry at the Institute of Biochemistry and Biophysics (Warsaw Poland), as previously described (12,74). Protein identities were confirmed by Western blotting using specific antibodies. Whole cell lysates and nuclear extracts were diluted by mixing with the same volume of a cytoplasmic buffer (40 mM HEPES pH 7.9, 150 mM NaCl, 4.5 mM MgCl_2_, 0.5 mM DTT), and used in the RNA binding assay, as described above.

### Protein expression

cDNAs encoding full length PTBP1 (short and long form), PTBP2 and hnRNP A1 were cloned into pET42a or pET28a vectors (Novagen) and the encoded proteins expressed in *E. coli* and purified on Ni-NTA agarose beads (Qiagen) using standard protocols. The proteins containing a 5xHistidine tag (pET28a) or both 5xHistidine and GST tags (pET42a) were used in binding assays without further purification. *In vitro* protein synthesis by the coupled transcription and translation (TnT) system (Promega) was carried out in the presence of ^35^S methionine in either wheat germ or rabbit reticulocyte lysates, as suggested by the manufacturer.

### Electrophoretic Mobility Shift Assay (EMSA)

The formation of RNA/protein complexes was monitored by Electrophoretic Mobility Shift Assay (EMSA) using RNA probes labeled at the 5’ end with ^32^P, as described previously (106). Typically, each sample in a total volume of 10 µl contained 5 µl of EMSA buffer (40 mM HEPES pH 7.9, 150 mM NaCl, 4.5 mM MgCl_2_, 0.5 mM DTT 0.2 µg/µl of yeast tRNA, 5% glycerol) and 5 µl of a TnT lysate, cytoplasmic extract, or recombinant protein (total 0.5-1 µg). The samples were incubated for 15 min on ice, resolved in a 6% native polyacrylamide gel and the radioactive RNA probe and its complexes detected by autoradiography.

## ACKNOWLEDGEMENTS

We thank J. Oledzki, A. Malinowska and M. Dadlez (Laboratory of Mass Spectrometry, Institute of Biochemistry and Biophysics, Polish Academy of Sciences) for mass spectrometry analysis. We also thank the following researchers for providing reagents and helpful suggestions: F. Conlon, K. Mason, J. Giudice and E. Blue (UNC, Chapel Hill), A. Mayeda (Fujita Health University), M. Nagiec (Broad Institute), and J. Deshaies and C. Vande Velde (University of Montreal). L. Tong (Columbia University) is greatly appreciated for critical reading of the manuscript. This study was funded by the NIH grant GM 29832.

## REFERENCES

1. Dominski, Z. and Marzluff, W.F. (2007) Formation of the 3’ end of histone mRNA: getting closer to the end. Gene, 396, 373–390.

2. Dominski, Z., Yang, X.C. and Marzluff, W.F. (2005) The Polyadenylation Factor CPSF-73 Is Involved in Histone-Pre-mRNA Processing. Cell, 123, 37–48.

3. Sun, Y., Zhang, Y., Aik, W.S., Yang, X.C., Marzluff, W.F., Walz, T., Dominski, Z. and Tong, L. (2020) Structure of an active human histone pre-mRNA 3’-end processing machinery. Science, 367, 700–703.

4. Dominski, Z. and Tong, L. (2021) U7 deciphered: the mechanism that forms the unusual 3’ end of metazoan replication-dependent histone mRNAs. Biochem. Soc. Trans., 49, 2229–2240.

5. Liu, H. and Moore, C.L. (2021) On the Cutting Edge: Regulation and Therapeutic Potential of the mRNA 3’ End Nuclease. Trends Biochem. Sci., 46, 772–784.

6. Mandel, C.R., Kaneko, S., Zhang, H., Gebauer, D., Vethantham, V., Manley, J.L. and Tong, L. (2006) Polyadenylation factor CPSF-73 is the pre-mRNA 3’-end-processing endonuclease. Nature, 444, 953–956.

7. Schaufele, F., Gilmartin, G.M., Bannwarth, W. and Birnstiel, M.L. (1986) Compensatory mutations suggest that base-pairing with a small nuclear RNA is required to form the 3’ end of H3 messenger RNA. Nature, 323, 777–781.

8. Bond, U.M., Yario, T.A. and Steitz, J.A. (1991) Multiple processing-defective mutations in a mammalian histone premessenger RNA are suppressed by compensatory changes in U7 RNA both in vivo and in vitro. Genes Dev, 5, 1709–1722.

9. Schumperli, D. and Pillai, R.S. (2004) The special Sm core structure of the U7 snRNP: far-reaching significance of a small nuclear ribonucleoprotein. Cell Mol. Life Sci, 61, 2560–2570.

10. Khusial, P., Plaag, R. and Zieve, G.W. (2005) LSm proteins form heptameric rings that bind to RNA via repeating motifs. Trends Biochem. Sci, 30, 522–528.

11. Yang, X.C., Xu, B., Sabath, I., Kunduru, L., Burch, B.D., Marzluff, W.F. and Dominski, Z. (2011) FLASH is required for the endonucleolytic cleavage of histone pre-mRNAs but is dispensable for the 5’ exonucleolytic degradation of the downstream cleavage product. Mol. Cell Biol, 31, 1492–1502.

12. Yang, X.C., Sabath, I., Debski, J., Kaus-Drobek, M., Dadlez, M., Marzluff, W.F. and Dominski, Z. (2013) A Complex Containing the CPSF73 Endonuclease and Other Polyadenylation Factors Associates with U7 snRNP and Is Recruited to Histone Pre-mRNA for 3’-End Processing. Molecular and Cellular Biology, 33, 28–37.

13. Battle, D.J., Kasim, M., Yong, J., Lotti, F., Lau, C.K., Mouaikel, J., Zhang, Z., Han, K., Wan, L. and Dreyfuss, G. (2006) The SMN complex: an assembly machine for RNPs. Cold Spring Harb. Symp. Quant. Biol., 71, 313–320.

14. Neuenkirchen, N., Chari, A. and Fischer, U. (2008) Deciphering the assembly pathway of Sm-class U snRNPs. FEBS Lett., 582, 1997–2003.

15. Gruss, O.J., Meduri, R., Schilling, M. and Fischer, U. (2017) UsnRNP biogenesis: mechanisms and regulation. Chromosoma, 126, 577–593.

16. Li, D.K., Tisdale, S., Lotti, F. and Pellizzoni, L. (2014) SMN control of RNP assembly: from post-transcriptional gene regulation to motor neuron disease. Semin Cell Dev Biol, 32, 22–29.

17. Meister, G. and Fischer, U. (2002) Assisted RNP assembly: SMN and PRMT5 complexes cooperate in the formation of spliceosomal UsnRNPs. EMBO J., 21, 5853–5863.

18. Raker, V.A., Hartmuth, K., Kastner, B. and Lührmann, R. (1999) Spliceosomal U snRNP core assembly: Sm proteins assemble onto an Sm site RNA nonanucleotide in a specific and thermodynamically stable manner. Molecular and Cellular Biology, 19, 6554–6565.

19. Paknia, E., Chari, A., Stark, H. and Fischer, U. (2016) The Ribosome Cooperates with the Assembly Chaperone pICln to Initiate Formation of snRNPs. Cell Rep, 16, 3103–3112.

20. Friesen, W.J., Paushkin, S., Wyce, A., Massenet, S., Pesiridis, G.S., Van Duyne, G., Rappsilber, J., Mann, M. and Dreyfuss, G. (2001) The methylosome, a 20S complex containing JBP1 and pICln, produces dimethylarginine-modified Sm proteins. Molecular and Cellular Biology, 21, 8289–8300.

21. Meister, G., Eggert, C., Buhler, D., Brahms, H., Kambach, C. and Fischer, U. (2001) Methylation of Sm proteins by a complex containing PRMT5 and the putative U snRNP assembly factor pICln. Curr. Biol., 11, 1990–1994.

22. Brahms, H., Meheus, L., de Brabandere, V., Fischer, U. and Luhrmann, R. (2001) Symmetrical dimethylation of arginine residues in spliceosomal Sm protein B/B’ and the Sm-like protein LSm4, and their interaction with the SMN protein. RNA, 7, 1531–1542.

23. Zhang, R., So, B.R., Li, P., Yong, J., Glisovic, T., Wan, L. and Dreyfuss, G. (2011) Structure of a key intermediate of the SMN complex reveals Gemin2’s crucial function in snRNP assembly. Cell, 146, 384–395.

24. Battle, D.J., Lau, C.K., Wan, L., Deng, H., Lotti, F. and Dreyfuss, G. (2006) The Gemin5 protein of the SMN complex identifies snRNAs. Mol. Cell, 23, 273–279.

25. Lau, C.K., Bachorik, J.L. and Dreyfuss, G. (2009) Gemin5-snRNA interaction reveals an RNA binding function for WD repeat domains. Nat. Struct. Mol. Biol., 16, 486–491.

26. Yong, J., Kasim, M., Bachorik, J.L., Wan, L. and Dreyfuss, G. (2010) Gemin5 delivers snRNA precursors to the SMN complex for snRNP biogenesis. Mol. Cell, 38, 551–562.

27. Friesen, W.J., Massenet, S., Paushkin, S., Wyce, A. and Dreyfuss, G. (2001) SMN, the product of the spinal muscular atrophy gene, binds preferentially to dimethylarginine-containing protein targets. Mol. Cell, 7, 1111–1117.

28. Cote, J. and Richard, S. (2005) Tudor domains bind symmetrical dimethylated arginines. J. Biol. Chem., 280, 28476–28483.

29. Selenko, P., Sprangers, R., Stier, G., Buhler, D., Fischer, U. and Sattler, M. (2001) SMN tudor domain structure and its interaction with the Sm proteins. Nat. Struct. Biol., 8, 27–31.

30. Tripsianes, K., Madl, T., Machyna, M., Fessas, D., Englbrecht, C., Fischer, U., Neugebauer, K.M. and Sattler, M. (2011) Structural basis for dimethylarginine recognition by the Tudor domains of human SMN and SPF30 proteins. Nat. Struct. Mol. Biol., 18, 1414–1420.

31. Lefebvre, S., Burglen, L., Reboullet, S., Clermont, O., Burlet, P., Viollet, L., Benichou, B., Cruaud, C., Millasseau, P., Zeviani, M. et al. (1995) Identification and characterization of a spinal muscular atrophy-determining gene. Cell, 80, 155–165.

32. Chari, A., Paknia, E. and Fischer, U. (2009) The role of RNP biogenesis in spinal muscular atrophy. Curr. Opin. Cell Biol., 21, 387–393.

33. Pellizzoni, L. (2007) Chaperoning ribonucleoprotein biogenesis in health and disease. EMBO Rep, 8, 340–345.

34. Iannaccone, S.T., Smith, S.A. and Simard, L.R. (2004) Spinal muscular atrophy. Curr. Neurol. Neurosci. Rep., 4, 74–80.

35. Gubitz, A.K., Feng, W. and Dreyfuss, G. (2004) The SMN complex. Exp. Cell Res, 296, 51–56.

36. Faravelli, I., Riboldi, G.M., Rinchetti, P. and Lotti, F. (2023) The SMN Complex at the Crossroad between RNA Metabolism and Neurodegeneration. Int. J. Mol. Sci., 24.

37. Zhang, Z., Lotti, F., Dittmar, K., Younis, I., Wan, L., Kasim, M. and Dreyfuss, G. (2008) SMN deficiency causes tissue-specific perturbations in the repertoire of snRNAs and widespread defects in splicing. Cell, 133, 585–600.

38. Winkler, C., Eggert, C., Gradl, D., Meister, G., Giegerich, M., Wedlich, D., Laggerbauer, B. and Fischer, U. (2005) Reduced U snRNP assembly causes motor axon degeneration in an animal model for spinal muscular atrophy. Genes Dev, 19, 2320–2330.

39. Lotti, F., Imlach, W.L., Saieva, L., Beck, E.S., Hao le, T., Li, D.K., Jiao, W., Mentis, G.Z., Beattie, C.E., McCabe, B.D. et al. (2012) An SMN-dependent U12 splicing event essential for motor circuit function. Cell, 151, 440–454.

40. Monani, U.R. (2005) Spinal muscular atrophy: a deficiency in a ubiquitous protein; a motor neuron-specific disease. Neuron, 48, 885–896.

41. Burghes, A.H. and Beattie, C.E. (2009) Spinal muscular atrophy: why do low levels of survival motor neuron protein make motor neurons sick? Nat. Rev. Neurosci., 10, 597–609.

42. Fallini, C., Bassell, G.J. and Rossoll, W. (2012) Spinal muscular atrophy: the role of SMN in axonal mRNA regulation. Brain Res., 1462, 81–92.

43. Pillai, R.S., Grimmler, M., Meister, G., Will, C.L., Luhrmann, R., Fischer, U. and Schumperli, D. (2003) Unique Sm core structure of U7 snRNPs: assembly by a specialized SMN complex and the role of a new component, Lsm11, in histone RNA processing. Genes Dev, 17, 2321–2333.

44. Yang, X.C., Desotell, A., Lin, M.H., Paige, A.S., Malinowska, A., Sun, Y., Aik, W.S., Dadlez, M., Tong, L. and Dominski, Z. (2023) In vitro methylation of the U7 snRNP subunits Lsm11 and SmE by the PRMT5/MEP50/pICln methylosome. RNA.

45. Tisdale, S., Lotti, F., Saieva, L., Van Meerbeke, J.P., Crawford, T.O., Sumner, C.J., Mentis, G.Z. and Pellizzoni, L. (2013) SMN is essential for the biogenesis of U7 small nuclear ribonucleoprotein and 3’-end formation of histone mRNAs. Cell Rep, 5, 1187–1195.

46. Tisdale, S., Van Alstyne, M., Simon, C.M., Mentis, G.Z. and Pellizzoni, L. (2022) SMN controls neuromuscular junction integrity through U7 snRNP. Cell Rep, 40, 111393.

47. Garcia-Blanco, M.A., Jamison, S.F. and Sharp, P.A. (1989) Identification and purification of a 62,000-dalton protein that binds specifically to the polypyrimidine tract of introns. Genes Dev, 3, 1874–1886.

48. Gil, A., Sharp, P.A., Jamison, S.F. and Garcia-Blanco, M.A. (1991) Characterization of cDNAs encoding the polypyrimidine tract-binding protein. Genes Dev, 5, 1224–1236.

49. Ghetti, A., Pinol-Roma, S., Michael, W.M., Morandi, C. and Dreyfuss, G. (1992) hnRNP I, the polypyrimidine tract-binding protein: distinct nuclear localization and association with hnRNAs. Nucleic Acids Res., 20, 3671–3678.

50. Oberstrass, F.C., Auweter, S.D., Erat, M., Hargous, Y., Henning, A., Wenter, P., Reymond, L., Amir-Ahmady, B., Pitsch, S., Black, D.L. et al. (2005) Structure of PTB bound to RNA: specific binding and implications for splicing regulation. Science, 309, 2054–2057.

51. Auweter, S.D. and Allain, F.H. (2008) Structure-function relationships of the polypyrimidine tract binding protein. Cell Mol Life Sci, 65, 516–527.

52. Clerte, C. and Hall, K.B. (2009) The domains of polypyrimidine tract binding protein have distinct RNA structural preferences. Biochemistry, 48, 2063–2074.

53. Mayeda, A., Munroe, S.H., Cáceres, J.F. and Krainer, A.R. (1994) Function of conserved domains of hnRNP A1 and other hnRNP A/B proteins. EMBO J., 13, 5483–5495.

54. Jean-Philippe, J., Paz, S. and Caputi, M. (2013) hnRNP A1: the Swiss army knife of gene expression. Int. J. Mol. Sci., 14, 18999–19024.

55. Beusch, I., Barraud, P., Moursy, A., Clery, A. and Allain, F.H. (2017) Tandem hnRNP A1 RNA recognition motifs act in concert to repress the splicing of survival motor neuron exon 7. Elife, 6.

56. Kashima, T. and Manley, J.L. (2003) A negative element in SMN2 exon 7 inhibits splicing in spinal muscular atrophy. Nat. Genet., 34, 460–463.

57. Coelho, M.B., Ascher, D.B., Gooding, C., Lang, E., Maude, H., Turner, D., Llorian, M., Pires, D.E., Attig, J. and Smith, C.W. (2016) Functional interactions between polypyrimidine tract binding protein and PRI peptide ligand containing proteins. Biochem. Soc. Trans., 44, 1058–1065.

58. Coelho, M.B., Attig, J., Bellora, N., Konig, J., Hallegger, M., Kayikci, M., Eyras, E., Ule, J. and Smith, C.W. (2015) Nuclear matrix protein Matrin3 regulates alternative splicing and forms overlapping regulatory networks with PTB. EMBO J., 34, 653–668.

59. Joshi, A., Coelho, M.B., Kotik-Kogan, O., Simpson, P.J., Matthews, S.J., Smith, C.W. and Curry, S. (2011) Crystallographic analysis of polypyrimidine tract-binding protein-Raver1 interactions involved in regulation of alternative splicing. Structure, 19, 1816–1825.

60. Henneberg, B., Swiniarski, S., Sabine, B. and Illenberger, S. (2010) A conserved peptide motif in Raver2 mediates its interaction with the polypyrimidine tract-binding protein. Exp. Cell Res., 316, 966–979.

61. Kleinhenz, B., Fabienke, M., Swiniarski, S., Wittenmayer, N., Kirsch, J., Jockusch, B.M., Arnold, H.H. and Illenberger, S. (2005) Raver2, a new member of the hnRNP family. FEBS Lett., 579, 4254–4258.

62. Coelho, M.B., Attig, J., Ule, J. and Smith, C.W. (2016) Matrin3: connecting gene expression with the nuclear matrix. Wiley Interdiscip Rev RNA, 7, 303–315.

63. Sawicka, K., Bushell, M., Spriggs, K.A. and Willis, A.E. (2008) Polypyrimidine-tract-binding protein: a multifunctional RNA-binding protein. Biochem. Soc. Trans., 36, 641–647.

64. Vuong, J.K., Lin, C.H., Zhang, M., Chen, L., Black, D.L. and Zheng, S. (2016) PTBP1 and PTBP2 Serve Both Specific and Redundant Functions in Neuronal Pre-mRNA Splicing. Cell Rep, 17, 2766–2775.

65. Spellman, R., Llorian, M. and Smith, C.W. (2007) Crossregulation and functional redundancy between the splicing regulator PTB and its paralogs nPTB and ROD1. Mol. Cell, 27, 420–434.

66. Ashiya, M. and Grabowski, P.J. (1997) A neuron-specific splicing switch mediated by an array of pre-mRNA repressor sites: evidence of a regulatory role for the polypyrimidine tract binding protein and a brain-specific PTB counterpart. RNA, 3, 996–1015.

67. Keppetipola, N., Sharma, S., Li, Q. and Black, D.L. (2012) Neuronal regulation of pre-mRNA splicing by polypyrimidine tract binding proteins, PTBP1 and PTBP2. Crit Rev Biochem Mol Biol, 47, 360–378.

68. Coutinho-Mansfield, G.C., Xue, Y., Zhang, Y. and Fu, X.D. (2007) PTB/nPTB switch: a post-transcriptional mechanism for programming neuronal differentiation. Genes Dev, 21, 1573–1577.

69. Boutz, P.L., Stoilov, P., Li, Q., Lin, C.H., Chawla, G., Ostrow, K., Shiue, L., Ares, M., Jr. and Black, D.L. (2007) A post-transcriptional regulatory switch in polypyrimidine tract-binding proteins reprograms alternative splicing in developing neurons. Genes Dev, 21, 1636–1652.

70. Cotten, M., Gick, O., Vasserot, A., Schaffner, G. and Birnstiel, M.L. (1988) Specific contacts between mammalian U7 snRNA and histone precursor RNA are indispensable for the in vitro RNA processing reaction. EMBO J., 7, 801–808.

71. Baillat, D., Hakimi, M.A., Naar, A.M., Shilatifard, A., Cooch, N. and Shiekhattar, R. (2005) Integrator, a multiprotein mediator of small nuclear RNA processing, associates with the C-terminal repeat of RNA polymerase II. Cell, 123, 265–276.

72. Egloff, S., O’Reilly, D. and Murphy, S. (2008) Expression of human snRNA genes from beginning to end. Biochem. Soc. Trans, 36, 590–594.

73. Matera, A.G., Terns, R.M. and Terns, M.P. (2007) Non-coding RNAs: lessons from the small nuclear and small nucleolar RNAs. Nat. Rev. Mol. Cell Biol, 8, 209–220.

74. Skrajna, A., Yang, X.C., Dadlez, M., Marzluff, W.F. and Dominski, Z. (2018) Protein composition of catalytically active U7-dependent processing complexes assembled on histone pre-mRNA containing biotin and a photo-cleavable linker. Nucleic Acids Res., 46, 4752–4770.

75. Skrajna, A., Yang, X.C. and Dominski, Z. (2019) Single-step Purification of Macromolecular Complexes Using RNA Attached to Biotin and a Photo-cleavable Linker. J Vis Exp.

76. Grimm, C., Stefanovic, B. and Schümperli, D. (1993) The low abundance of U7 snRNA is partly determined by its Sm binding site. EMBO J., 12, 1229–1238.

77. Stefanovic, B., Hackl, W., Lührmann, R. and Schümperli, D. (1995) Assembly, nuclear import and function of U7 snRNPs studied by microinjection of synthetic U7 RNA into *Xenopus* oocytes. Nucleic Acids Res., 23, 3141–3151.

78. Singh, R., Valcárcel, J. and Green, M.R. (1995) Distinct binding specificities and functions of higher eukaryotic polypyrimidine tract-binding proteins. Science, 268, 1173–1176.

79. Perez, I., Lin, C.H., McAfee, J.G. and Patton, J.G. (1997) Mutation of PTB binding sites causes misregulation of alternative 3’ splice site selection in vivo. RNA, 3, 764–778.

80. Monie, T.P., Hernandez, H., Robinson, C.V., Simpson, P., Matthews, S. and Curry, S. (2005) The polypyrimidine tract binding protein is a monomer. RNA, 11, 1803–1808.

81. Clerte, C. and Hall, K.B. (2006) Characterization of multimeric complexes formed by the human PTB1 protein on RNA. RNA, 12, 457–475.

82. Yaniv, K. and Yisraeli, J.K. (2002) The involvement of a conserved family of RNA binding proteins in embryonic development and carcinogenesis. Gene, 287, 49–54.

83. Bell, J.L., Wachter, K., Muhleck, B., Pazaitis, N., Kohn, M., Lederer, M. and Huttelmaier, S. (2013) Insulin-like growth factor 2 mRNA-binding proteins (IGF2BPs): post-transcriptional drivers of cancer progression? Cell Mol Life Sci, 70, 2657–2675.

84. Yisraeli, J.K. (2005) VICKZ proteins: a multi-talented family of regulatory RNA-binding proteins. Biol Cell, 97, 87–96.

85. Kovalevich, J., Santerre, M. and Langford, D. (2021) Considerations for the Use of SH-SY5Y Neuroblastoma Cells in Neurobiology. Methods Mol. Biol., 2311, 9–23.

86. Xu, C., Ishikawa, H., Izumikawa, K., Li, L., He, H., Nobe, Y., Yamauchi, Y., Shahjee, H.M., Wu, X.H., Yu, Y.T. et al. (2016) Structural insights into Gemin5-guided selection of pre-snRNAs for snRNP assembly. Genes Dev, 30, 2376–2390.

87. Jin, W., Wang, Y., Liu, C.P., Yang, N., Jin, M., Cong, Y., Wang, M. and Xu, R.M. (2016) Structural basis for snRNA recognition by the double-WD40 repeat domain of Gemin5. Genes Dev, 30, 2391–2403.

88. Tang, X., Bharath, S.R., Piao, S., Tan, V.Q., Bowler, M.W. and Song, H. (2016) Structural basis for specific recognition of pre-snRNA by Gemin5. Cell Res., 26, 1353–1356.

89. Wahl, M.C. and Fischer, U. (2016) The right pick: structural basis of snRNA selection by Gemin5. Genes Dev, 30, 2341–2344.

90. Burd, C.G. and Dreyfuss, G. (1994) RNA binding specificity of hnRNP A1: Significance of hnRNP A1 high-affinity binding sites in pre-mRNA splicing. EMBO J., 13, 1197–1204.

91. Raker, V.A., Plessel, G. and Lührmann, R. (1996) The snRNP core assembly pathway: Identification of stable core protein heteromeric complexes and an snRNP subcore particle *in vitro*. EMBO J., 15, 2256–2269.

92. Urlaub, H., Raker, V.A., Kostka, S. and Lührmann, R. (2001) Sm protein-Sm site RNA interactions within the inner ring of the spliceosomal snRNP core structure. EMBO J., 20, 187–196.

93. Will, C.L. and Luhrmann, R. (2001) Spliceosomal UsnRNP biogenesis, structure and function. Curr. Opin. Cell Biol, 13, 290–301.

94. Perez, I., McAfee, J.G. and Patton, J.G. (1997) Multiple RRMs contribute to RNA binding specificity and affinity for polypyrimidine tract binding protein. Biochemistry, 36, 11881–11890.

95. Marz, M., Mosig, A., Stadler, B.M. and Stadler, P.F. (2007) U7 snRNAs: a computational survey. Genomics Proteomics Bioinformatics, 5, 187–195.

96. Lührmann, R., Kastner, B. and Bach, M. (1990) Structure of spliceosomal snRNPs and their role in pre-mRNA splicing. Biochim. Biophys. Acta Gene Struct. Expression, 1087, 265–292.

97. Bucholc, K., Aik, W.S., Yang, X.C., Wang, K., Zhou, Z.H., Dadlez, M., Marzluff, W.F., Tong, L. and Dominski, Z. (2020) Composition and processing activity of a semi-recombinant holo U7 snRNP. Nucleic Acids Res., 48, 1508–1530.

98. Yang, X.C., Sun, Y., Aik, W.S., Marzluff, W.F., Tong, L. and Dominski, Z. (2020) Studies with recombinant U7 snRNP demonstrate that CPSF73 is both an endonuclease and a 5’-3’ exonuclease. RNA, 26, 1345–1359.

99. Dai, S., Wang, C., Zhang, C., Feng, L., Zhang, W., Zhou, X., He, Y., Xia, X., Chen, B. and Song, W. (2022) PTB: Not just a polypyrimidine tract-binding protein. J. Cell. Physiol., 237, 2357–2373.

100. Hu, J., Qian, H., Xue, Y. and Fu, X.D. (2018) PTB/nPTB: master regulators of neuronal fate in mammals. Biophys Rep, 4, 204–214.

101. Vitali, F., Henning, A., Oberstrass, F.C., Hargous, Y., Auweter, S.D., Erat, M. and Allain, F.H. (2006) Structure of the two most C-terminal RNA recognition motifs of PTB using segmental isotope labeling. EMBO J., 25, 150–162.

102. Lamichhane, R., Daubner, G.M., Thomas-Crusells, J., Auweter, S.D., Manatschal, C., Austin, K.S., Valniuk, O., Allain, F.H. and Rueda, D. (2010) RNA looping by PTB: Evidence using FRET and NMR spectroscopy for a role in splicing repression. Proc. Natl. Acad. Sci. U. S. A., 107, 4105–4110.

103. Jonson, L., Vikesaa, J., Krogh, A., Nielsen, L.K., Hansen, T., Borup, R., Johnsen, A.H., Christiansen, J. and Nielsen, F.C. (2007) Molecular composition of IMP1 ribonucleoprotein granules. Mol Cell Proteomics, 6, 798–811.

104. Shamoo, Y., Krueger, U., Rice, L.M., Williams, K.R. and Steitz, T.A. (1997) Crystal structure of the two RNA binding domains of human hnRNP A1 at 1.75 A resolution. Nat. Struct. Biol., 4, 215–222.

105. Dominski, Z., Yang, X.C., Kaygun, H., Dadlez, M. and Marzluff, W.F. (2003) A 3’ exonuclease that specifically interacts with the 3’ end of histone mRNA. Mol. Cell, 12, 295–305.

106. Yang, X.C., Purdy, M., Marzluff, W.F. and Dominski, Z. (2006) Characterization of 3’hExo, a 3’ exonuclease specifically interacting with the 3’ end of histone mRNA. J. Biol. Chem., 281, 30447–30454.

107. Tan, D., Marzluff, W.F., Dominski, Z. and Tong, L. (2013) Structure of histone mRNA stem-loop, human stem-loop binding protein, and 3’hExo ternary complex. Science, 339, 318–321.

108. Hoefig, K.P. and Heissmeyer, V. (2014) Degradation of oligouridylated histone mRNAs: see UUUUU and goodbye. Wiley Interdiscip Rev RNA, 5, 577–589.

109. Hoefig, K.P., Rath, N., Heinz, G.A., Wolf, C., Dameris, J., Schepers, A., Kremmer, E., Ansel, K.M. and Heissmeyer, V. (2013) Eri1 degrades the stem-loop of oligouridylated histone mRNAs to induce replication-dependent decay. Nat. Struct. Mol. Biol, 20, 73–81.

110. Kolev, N.G. and Steitz, J.A. (2006) In vivo assembly of functional U7 snRNP requires RNA backbone flexibility within the Sm-binding site. Nat. Struct. Mol. Biol, 13, 347–353.

111. Rossoll, W., Kroning, A.K., Ohndorf, U.M., Steegborn, C., Jablonka, S. and Sendtner, M. (2002) Specific interaction of Smn, the spinal muscular atrophy determining gene product, with hnRNP-R and gry-rbp/hnRNP-Q: a role for Smn in RNA processing in motor axons? Hum. Mol. Genet, 11, 93–105.

112. Gromak, N., Rideau, A., Southby, J., Scadden, A.D., Gooding, C., Huttelmaier, S., Singer, R.H. and Smith, C.W. (2003) The PTB interacting protein raver1 regulates alpha-tropomyosin alternative splicing. EMBO J., 22, 6356–6364.

113. Rideau, A.P., Gooding, C., Simpson, P.J., Monie, T.P., Lorenz, M., Huttelmaier, S., Singer, R.H., Matthews, S., Curry, S. and Smith, C.W. (2006) A peptide motif in Raver1 mediates splicing repression by interaction with the PTB RRM2 domain. Nat. Struct. Mol. Biol., 13, 839–848.

114. Spellman, R., Rideau, A., Matlin, A., Gooding, C., Robinson, F., McGlincy, N., Grellscheid, S.N., Southby, J., Wollerton, M. and Smith, C.W. (2005) Regulation of alternative splicing by PTB and associated factors. Biochem. Soc. Trans., 33, 457–460.

115. Kafasla, P., Morgner, N., Poyry, T.A., Curry, S., Robinson, C.V. and Jackson, R.J. (2009) Polypyrimidine tract binding protein stabilizes the encephalomyocarditis virus IRES structure via binding multiple sites in a unique orientation. Mol. Cell, 34, 556–568.

116. Huttelmaier, S., Illenberger, S., Grosheva, I., Rudiger, M., Singer, R.H. and Jockusch, B.M. (2001) Raver1, a dual compartment protein, is a ligand for PTB/hnRNPI and microfilament attachment proteins. J. Cell Biol., 155, 775–786.

117. Zieseniss, A., Schroeder, U., Buchmeier, S., Schoenenberger, C.A., van den Heuvel, J., Jockusch, B.M. and Illenberger, S. (2007) Raver1 is an integral component of muscle contractile elements. Cell Tissue Res, 327, 583–594.

118. Lee, J.H., Rangarajan, E.S., Yogesha, S.D. and Izard, T. (2009) Raver1 interactions with vinculin and RNA suggest a feed-forward pathway in directing mRNA to focal adhesions. Structure, 17, 833–842.

119. Babic, I., Sharma, S. and Black, D.L. (2009) A role for polypyrimidine tract binding protein in the establishment of focal adhesions. Mol Cell Biol, 29, 5564–5577.

120. Dominski, Z., Sumerel, J., Hanson, R.J. and Marzluff, W.F. (1995) The polyribosomal protein bound to the 3’ end of histone mRNA can function in histone pre-mRNA processing. RNA, 1, 915–923.

121. Sun, Y., Aik, W.S., Yang, X.C., Marzluff, W.F., Dominski, Z. and Tong, L. (2021) Reconstitution and biochemical assays of an active human histone pre-mRNA 3’-end processing machinery. Methods Enzymol., 655, 291–324.

122. Mayeda, A. and Krainer, A.R. (1999) Preparation of HeLa cell nuclear and cytosolic S100 extracts for in vitro splicing. Methods Mol. Biol., 118, 309–314.

123. Sabath, I., Skrajna, A., Yang, X.C., Dadlez, M., Marzluff, W.F. and Dominski, Z. (2013) 3’-End processing of histone pre-mRNAs in Drosophila: U7 snRNP is associated with FLASH and polyadenylation factors. RNA, 19, 1726–1744.

